# Direct visualization of emergent metastatic features within an *ex vivo* model of the tumor microenvironment

**DOI:** 10.1101/2023.01.09.523294

**Authors:** Libi Anandi, Jeremy Garcia, Manon Ros, Libuše Janská, Josephine Liu, Carlos Carmona-Fontaine

**Affiliations:** Center for Genomics & Systems Biology, Department of Biology, New York University, New York, NY 10003, USA

## Abstract

Metabolic conditions such as hypoxia, nutrient starvation, and media acidification, together with interactions with stromal cells are critical drivers of metastasis. Since these conditions arise deep within tumor tissues with poor access to the bloodstream, the observation of nascent metastases *in vivo* is exceedingly challenging. On the other hand, conventional cell culture studies cannot capture the complex nature of metastatic processes. We thus designed and implemented an *ex vivo* model of the tumor microenvironment to study the emergence of metastatic features in tumor cells in their native 3-dimensional (3D) context. In this system, named 3MIC, tumor cells spontaneously create ischemic-like conditions, and it allows the direct visualization of tumor-stroma interactions with high spatial and temporal resolution. We studied how 3D tumor spheroids evolve in the 3MIC when cultured under different metabolic environments and in the presence or absence of stromal cells. Consistent with previous experimental and clinical data, we show that ischemic environments increase cell migration and invasion. Importantly, the 3MIC allowed us to directly observe the emergence of these pro-metastatic features with single-cell resolution allowing us to track how changes in tumor motility were modulated by macrophages and endothelial cells. With these tools, we determined that the acidification of the extracellular media was more important than hypoxia in the induction of pro-metastatic tumor features. We also illustrate how the 3MIC can be used to test the effects of anti-metastatic drugs on cells experiencing different metabolic conditions. Overall, the 3MIC allows us to directly observe the emergence of metastatic tumor features in a physiologically relevant model of the tumor microenvironment. This simple and cost-effective system can dissect the complexity of the tumor microenvironment to test perturbations that may prevent tumors from becoming metastatic.

## INTRODUCTION

Most cancer fatalities are directly or indirectly caused by metastases^1^. Thus, treating pre-metastatic tumor cells before they acquire migratory and invasive properties could dramatically reduce cancer mortality^2^. Recent studies using *in vivo* mouse models have provided novel insights into the metastatic process such as the role played by DNA damage through the promotion of the cGAS-STING pathway^3^, the increased metastatic efficiency of cell groups over single cells^4^, key roles of stromal and immune cells^5–7^, and conditions that facilitate the reactivation of dormant metastases and their engraftment into new tissues^8^.

The direct observation of nascent metastases remains elusive. Unfortunately, the observation of metastases *in vivo* requires sophisticated microscopy with costs that are prohibitively expensive for most laboratories^9–11^. Tools including quantification of circulating tumor cells, histological analyses of biopsies, and warm autopsies^12–18^ provide important information about late metastatic steps. However, early metastases are much more challenging to detect and to study^19–21^. This challenge is due in part to the complexity of cellular and molecular factors that regulate the emergence of metastasis ^22–25^. In addition, the initiation of metastasis is a stochastic process and thus when and where a metastatic clone emerges is unpredictable^26–28^. Amid these limitations, we sought to model the initial conditions that favor the emergence of metastasis *ex vivo*.

A model of a premetastatic tumor must recreate microenvironmental factors that are critical in the regulation of metastasis such as altered metabolic conditions^22–25^, and interactions with immune and stromal cells^22,29^. For example, in vivo imaging, molecular, and histological evidence have demonstrated and active role for tumor-infiltrated macrophages^5,30–32^ and fibroblasts^33–36^ in promoting and facilitating cancer invasion and metastasis. Metabolic stress in the tumor microenvironment is also associated with metastasis promotion^37–44^. Hypoxia is a common feature of solid tumors, and its pro-metastatic roles are well-established^40,41,45^. However, hypoxia rarely – if ever – occurs alone: as nutrients and oxygen diffuse into the tumor mass, oxygen and nutrients in the microenvironment become progressively scarcer while metabolic by-products, such as lactic acid, accumulate^46–53^. Most likely, multiple conditions within an ischemic microenvironment, including redox stress^54^, acidosis^43,55^, nutrient starvation^56^ rather than hypoxia alone, drive the initiation of metastasis.

By definition, ischemic conditions usually arise deep within tumors, and thus accessing and observing emergent metastases *in vivo* has been virtually impossible. Some aspects of tumor biology are well captured by *ex vivo* cultures such as organoids^57–64^ and other 3D tumor model systems^65–68^. Still, in these models, ischemic tumor cells remain buried within these structures and thus imaging tumor-stromal interactions in those regions poses an almost insurmountable challenge.

Here we present the development of an *ex vivo* model of the tumor microenvironment designed to visualize the transition of poorly motile primary tumor cells into migratory metastatic-like cells. This 3D Microenvironmental Chamber or 3MIC, models key tumor features including the infiltration of immune cells and the spontaneous formation of metabolic gradients that mimic the metabolic conditions within tumors. Due to its unique geometry, the 3MIC easily allows imaging ischemic cells with unprecedented temporal and spatial resolution. Using this system, we show that ischemic-like environments directly drive the emergence of metastatic features including increased cell migration, degradation of the extracellular matrix (ECM), and loss of epithelial features. Combining *in vivo* experiments with 3MIC cultures, we showed that these changes were reversible suggesting that metastatic features can arise even in the absence of clonal selection by hypoxia or other environmental challenges. We also show that tumor interactions with stromal cells such as macrophages and endothelial cells, increase the pro-metastatic effects of ischemia. Finally, we illustrate how the 3MIC can be used to test how local metabolic conditions may affect drug responses. In all, the 3MIC is an affordable cell culture system where different components of the tumor microenvironment can be carefully dissected. Its amenability for live imaging can complement *in vivo* studies to better understand and treat the emergence of metastases.

## RESULTS

### A 3D *ex vivo* model of the tumor microenvironment for the direct visualization of ischemic cells

Insufficient vascularization and excessive cell growth leave large ischemic regions within solid tumors^46–53^. Hypoxia, acidosis, and other metabolic stressors associated with ischemia, are known drivers of metastasis. Since these metabolic conditions often arise deep within tumors, visualizing the emergence of metastatic properties *in vivo* or in 3D organoids, presents unique technical challenges (Fig. 1A). To overcome these limitations, we first thought to use the **Me**tabolic **Mi**croenvironment **C**hamber (MEMIC), an *ex vivo* model of the tumor microenvironment that we developed to study the impact of ischemic stress on cell cultures^69–71^. In this system, cell monolayers form reproducible gradients of ischemia and by design imaging ischemic cells is as easy to as imaging well-nurtured cells.

**Figure 1:**
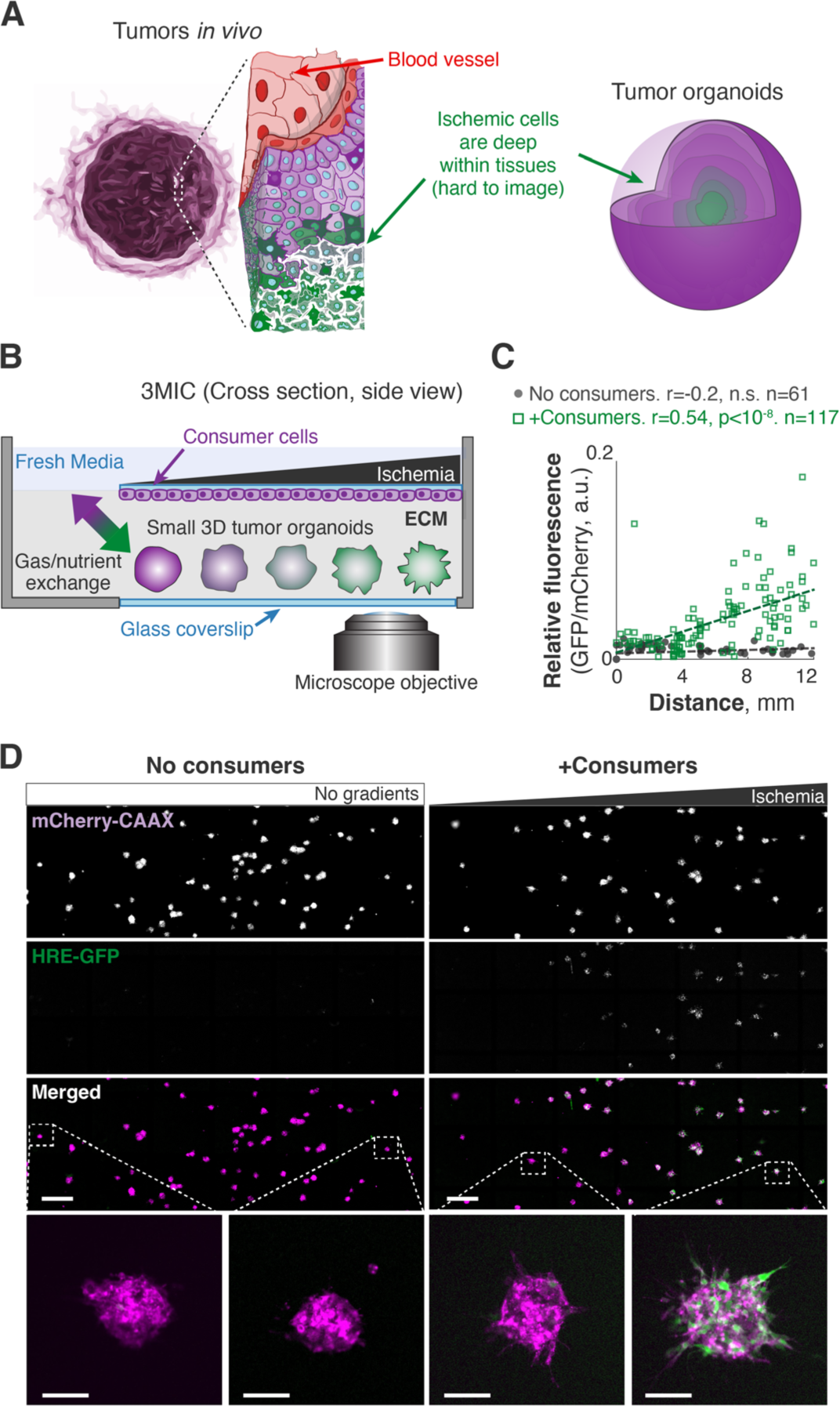
Design and implementation of a metabolic microenvironment chamber for 3D cultures. **A.** Sketch showing ischemic regions in solid tumor and conventional tumor organoids. Cells in these regions are hard to access and to image. **B.** Schematic cross-section of the 3MIC where a monolayer of “consumer” cells grows on the top of a small chamber. Tumor spheroids grow embedded in ECM within the same chamber and underneath consumer cells. This small chamber is connected to a large reservoir of fresh media from only one side (left) where nutrient and gas exchanges occur. **C.** Quantification of *per* spheroid GFP levels from HRE-GFP Lung KP 3D spheroids cultured in the 3MIC (∼100cells/spheroid) with or without consumer cells. GFP levels serve as a proxy for hypoxia and are normalized to constitutive mCherry-CAAX signal. Dashed line: linear fit. Data from a representative example of four biological replicates. **D.** Low magnification showing GFP and mCherry fluorescence in tumor spheroids cultured alone (left column) or in the presence of consumer cells (right column) along the 3MIC gradient. Bottom panels zoom into representative individual tumor spheroids. Bars: 1000µm, 100µm for insets.

Key features of metastatic cells involve morphological changes that are better modeled by 3D tumor structures than by cell monolayers. The MEMIC was designed for 2D cultures, and the density of these cells is crucial for the formation of metabolic gradients^71^. Unfortunately, this system did not allow achieving enough cell density in 3D cultures to form detectable ischemic gradients (Fig. S1A). We thus designed and developed a new system – which we named **3**D **mi**croenvironment **c**hamber (3MIC) – to study the impacts of the metabolic microenvironment on 3D cultures. The principle behind the 3MIC is similar to the one of the MEMIC: we grow a dense monolayer of cells inside a small chamber that is restricted from accessing nutrients and oxygen from all sides but one. The opening on this side connects to a large volume of fresh media and thus it acts as a source of nutrients and oxygen, while cells inside the chamber act as resource sinks. In the 3MIC, however, these cells are not the focus of our analyses and only act as nutrient consumers. These “consumer cells” grow upside-down on a coverslip at the top of the chamber and underneath them, we can introduce 3D tumor structures embedded in ECM. This two-tier system should allow us to have a high-density consumer cell layer producing strong metabolic gradients within a microenvironment shared by much sparser 3D tumor structures that in turn have a negligible impact on the metabolic conditions of the chamber (Fig. 1B; Fig. S1B). The location of consumer cells on top of the chamber allows an unobstructed view of the 3D tumor structures which will be closer to the objective of an inverted microscope (Fig. 1B). These experiments can use different combinations of cell types to form spheroids and the consumer layer. For simplicity, however, and unless mentioned otherwise, experiments shown here use the same cell type for both roles.

To test the 3MIC, we first confirmed that consumer cells form metabolic gradients. As a proxy for resource deprivation, we used a fluorescent hypoxia sensor (HRE-GFP^71^) which we expressed in a panel of epithelial tumor cells co-expressing constitutive plasma membrane fluorescence (mCherry-CAAX^71^). As in our MEMIC experiments, 2D monolayers of consumer cells formed strong ischemic gradients. For example, spheroids formed by Lung KP cells (derived from a murine model of lung adenocarcinoma driven by KRAS^G12D^ and loss of TP53^72^) cells displayed a strong gradient of GFP that increased with the distance to the opening of the 3MIC (Fig. S1C). We then aggregated cells into small tumor spheroids that we embedded in a collagen- and laminin-rich ECM scaffold and introduced them into 3MICs with or without monolayers of consumer cells.

As expected, spheroids in control 3MICs without consumer cells did not show signs of hypoxia (Fig. 1C,D). In contrast, tumor spheroids growing in 3MICs under consumer cells showed levels of the GFP hypoxia reporter that increased with the distance to the opening of the chamber (Fig. 1C,D). We obtained similar results with three independently derived clones of Lung KP cells and in experiments using different cell lines, and in different cell types expressing a different hypoxia reporter^73^ (Fig. S1D). These results show that consumer cells in the 3MIC form metabolic gradients that influence the local metabolic microenvironment of neighboring tumor spheroids. As these metabolic gradients in the 3MIC are perpendicular to our imaging path, we can observe how ischemic conditions affect tumor structures with unprecedented spatial and temporal resolution.

### Hypoxia is required but not sufficient to increase invasive features in tumor spheroids

In these experiments we observed that the morphology of tumor spheroids changed dramatically along the 3MIC. Ischemic spheroids acquired a ruffled and protrusive morphology suggesting an increased invasive behavior (Fig. 2A; Video S1). In contrast, well-nurtured spheroids close to the opening had a modest size increase and remained mostly smooth and rounded (Fig. 2A; Video S1). We decided to take advantage of the unique features of the 3MIC to identify the specific conditions that trigger these seemingly pro-metastatic features on ischemic spheroids.

**Figure 2:**
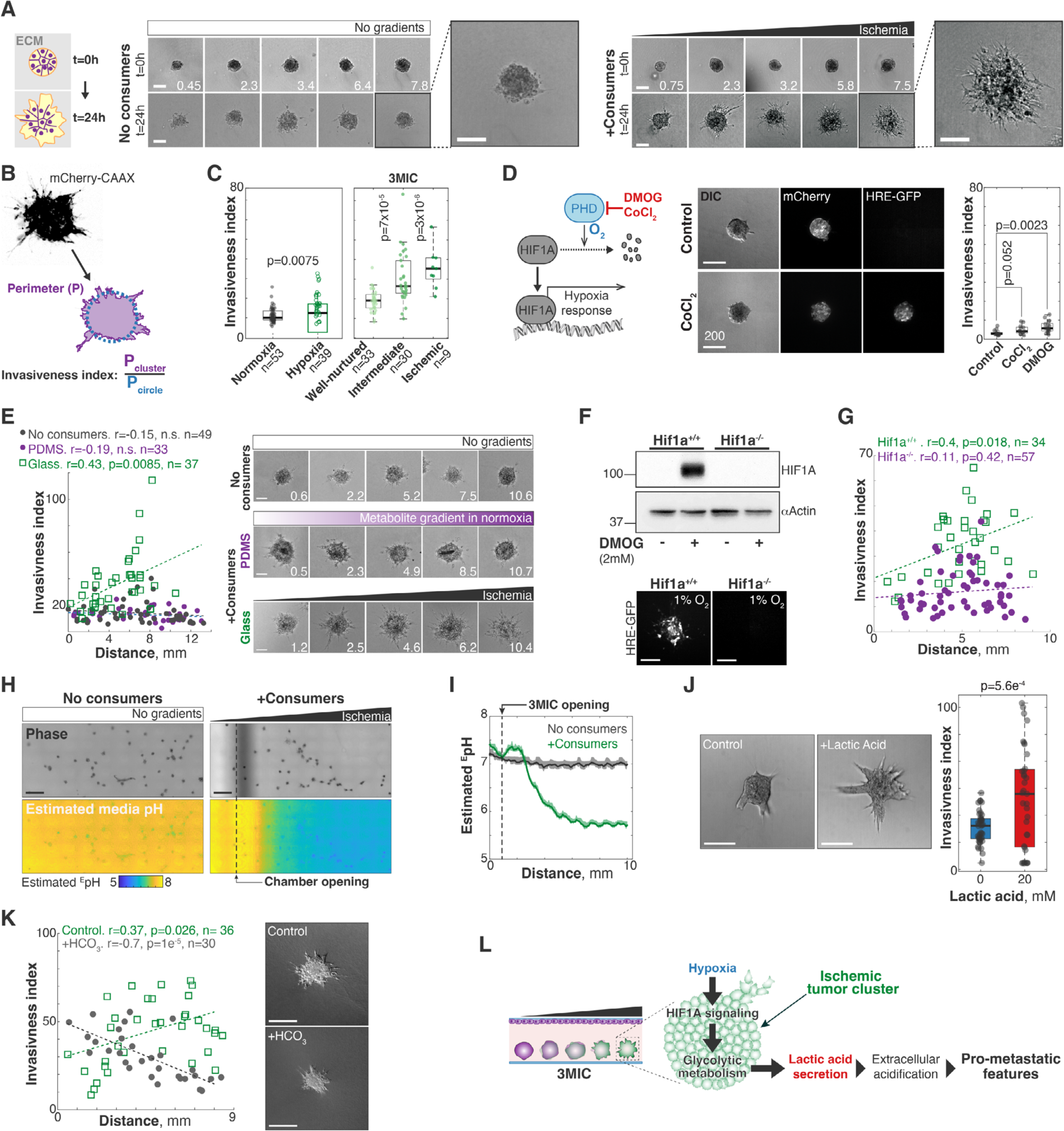
Low pH is responsible for the increased invasion of ischemic clusters in a HIF1A-dependent manner. **A.** Left: Lung KP spheroids imaged after 0h (top) and 24h (bottom) of growth in different regions of the 3MIC without consumer cells. These clusters grew uniformly and retained smooth and round shapes. Right: Similar data but for spheroids grown in the presence of consumer cells. Ischemic spheroids grew larger and more protrusive. Numbers in lower right corner denote the distance to the opening in mm. Bars: 100*μ*m. **B.** We defined invasiveness index, as the perimeter of tumor spheroid over the perimeter of a circle with the same area (1/circularity). **C.** Using the invasiveness index, we quantified the invasiveness of lung KP spheroids under different environments. Well-nurtured, intermediate, and ischemic conditions correspond to different regions of the 3MIC. Data points: *per* spheroid invasiveness. Data from a representative example of two biological replicates. **D.** Representative images and invasiveness index of untreated tumor spheroids compared to spheroids treated with CoCl_2_ or DMOG. Bars: 200µm. See supplementary figure 2 for images of DMOG-treated spheroids. **E.** Representative images of individual spheroids grown for 24h in the absence of consumer cells (top), or in the presence of consumer cells in 3MICs enclosed in PDMS (Polydimethylsiloxane) or glass (center and bottom, respectively). Numbers in lower right corner denote the distance to the chamber’s opening in mm. Bars: 100µm. Plots on the right show a quantification of these changes. Data points: invasiveness index of individual spheroids. Dash lines: linear fit. Data from a representative experiment of two biological replicates. **F.** Top: Anti-HIF1A blots of Hif1a^+/+^ and Hif1a^-/-^ cells treated or untreated with DMOG. Bottom: representative images of Hif1a^+/+^ and Hif1a^-/-^ HRE-GFP tumor spheroids cultured in a hypoxic incubator. **G.** Quantification of the invasiveness index of Hif1a^+/+^ and Hif1a^-/-^ spheroids along the 3MIC. Bars: 200µm. **H.** Representative ratiometric images used to estimate the extracellular pH (^E^pH) within 3MICs with or without consumer cells. **I.** Estimated extracellular pH over distance from the opening in 3MICs with or without consumer cells. **J.** Representative images and quantification of the invasiveness index of lung KP spheroids treated with 20mM of lactic acid. Bars: 200µm. **K.** Representative images and quantification of the invasiveness index of lung KP spheroids within 3MICs treated with 25mM of bicarbonate. Bars: 200µm. **L.** Model suggested by our data where increased invasion under ischemia is driven by low extracellular pH through a HIF1A-dependent mechanism.

There is ample evidence linking hypoxia to pro-metastatic tumor features^37–44^. However, whether additional metabolic stressors – such as nutrient deprivation or lactic acid accumulation – may contribute to the morphological changes we observe in the 3MIC. To rapidly compare the effects of different treatments on spheroid morphology, we defined *invasiveness index* as the inverse of their circularity. This index increases in cellular structures with irregular and protrusive shapes and remains small in smooth and homogeneous tumor spheroids (Fig. 2B).

We first tested whether hypoxia was sufficient to trigger morphological changes and increase cell migration in tumor spheroids. We compared spheroids cultured in regular petri dishes incubated under different oxygen tensions (21% and 1%) with spheroids cultured in the 3MIC. Tumor spheroids cultured in a hypoxic incubator increased levels of the GFP-based hypoxia sensor to levels comparable to ischemic clusters in the 3MIC (Fig. S2A). Hypoxic spheroids, however, were not as invasive as spheroids in the 3MIC and were only slightly more invasive than spheroids cultured in regular incubators (Fig. 2C). In similar experiments, we treated spheroids with dimethyloxalyglycine (DMOG) or cobalt chloride (CoCl_2_) – two prolyl hydroxylation inhibitors that stabilize the transcription factor Hypoxia-inducible factor 1-alpha (HIF1A) and trigger a hypoxic-like transcriptional response under normal oxygen tension^74^. As shown in Figure 2D, these spheroids showed only modest increases in invasion despite a strong activation of HIF1A response, as denoted by the strong increase in the GFP signal from the hypoxia reporter (Fig. 2D, Fig. S2B). These results show that hypoxia alone triggers relatively minor morphological changes and cannot fully recapitulate the changes that we observe in ischemic regions of the 3MIC.

Additional experiments however revealed that while not *sufficient*, hypoxia was *required* in altering spheroid morphologies in ischemia. First, we took advantage of the selective permeability of polydimethylsiloxane (PDMS)^69–71^. PDMS is permeable to oxygen and other gases, but not to soluble metabolites^75^. We thus constructed 3MICs using a PDMS membrane molded into the shape of a glass coverslip. In these modified 3MICs, cells form gradients of nutrients and metabolic byproducts, but oxygen rapidly equilibrates across the PDMS membrane and thus there is no hypoxia (Fig. S2C). Under these conditions, tumor spheroids did not show major morphological challenges, regardless of whether consumer cells were present or not (Fig. 2E; Fig. S2C). Second, we established that the changes displayed by ischemic spheroids require HIF1A signaling. Using CRISPR/Cas-9 we generated isogenic knockout Lung KP cells (Hif1a^-/-^). As expected, Hif1a^-/-^ cells did not have detectable levels of HIF1A-even after treatment with DMOG (Fig. 2F) and did increase their hypoxia-driven GFP levels when cultured in a hypoxic incubator (Fig. S2D). As in previous experiments, isogenic wildtype cells (Hif1a^+/+^) showed strong morphological changes along the 3MIC but these changes were severely impaired in spheroids formed by Hif1a^-/-^ cells (Fig. 2G). Altogether, these results show that hypoxia and HIF1A signaling are required but not sufficient for the morphological changes seen in ischemic clusters. As the effects of these perturbations alone are weak, we thought that cells need additional metabolic cues to fully adopt the morphologies we observe in ischemic spheroids.

### Media acidification is critical for the increase in metastatic features displayed by ischemic tumor cells

We then sought for additional factors that may increase the invasive capacity of tumor spheroids. A simple look with the naked eye at a 3MICs culture would suggest that strong changes in pH are occurring inside. We regularly use cell culture media with the pH indicator phenol red which show its typical red color near the opening of the chamber gradually yellowing deeper with the 3MIC (Fig. S2E). A ratiometric analysis of this change showed that extracellular pH levels drastically decrease in ischemic regions while they remained relatively constant in 3MICs without consumer cells (Fig. 2H,I).

Changes in intracellular pH were much milder (Fig. S2F). We thus focused on factors affecting media acidification such as lactic acid, which is abundantly produced and secreted by the glycolytic metabolism of tumors. Surprisingly, the simple addition of lactic acid to tumor spheroids under otherwise well-perfused and normal conditions, strongly increased their invasiveness (Fig. 2J). Consistent with a key role of low pH in promoting cell invasion, addition of bicarbonate to increase the buffering capacity of the media, eliminated the invasiveness of ischemic spheroids in the 3MIC (Fig. 2K). Importantly, acidification of the extracellular environment has been linked to increased tumor invasion and metastasis^43,76,77^.

Together, these results show that HIF1A signaling is required to increase the invasion of ischemic spheroids while a low pH is sufficient to do so. We speculate that these effects are linked as HIF1A increases lactic acid secretion through its regulation of glycolysis^78^. We think that in normal cultures, or in well-perfused tissues, lactic acid levels do not rise enough to produce a strong pH decrease. But in the 3MIC or in poorly vascularized tissues such as solid tumors, lactic acid is not cleared rapidly enough and thus it accumulates increasing cell invasion through a pH reduction (Fig. 2L). These experiments illustrate the importance of considering ischemia as a whole rather than hypoxia alone and they illustrate the ease of modeling these complex metabolic microenvironments in the 3MIC.

### Ischemia stimulates persistent cell migration

The increased invasion of ischemic spheroids was conserved across a panel of human and mouse cell lines (Fig. S3A,B). To better understand the mechanisms behind these changes, we first ruled out a major contribution for cell proliferation changes that can also drive the expansion of cell populations^79,80^. In fact, only a small number of cells in our 3D cultures were actively proliferating as revealed by a sparse signal for phospho-H3 immunostaining (Fig. S3C) and by FUCCI-based live imaging of the cell cycle (Fig. S3D; Video S2).

In contrast, live imaging analyses showed a significantly higher cell dispersion in ischemic clusters quantified as a decrease in local cell density (Fig. 3A,B; Video S3)^69^. Taking advantage of the high temporal and spatial resolution in live microscopy afforded by the 3MIC, we measured the migratory properties of individual tumor cells. Cells in these 3D structures are tightly packed muddling cell detection and tracking. To avoid this problem, we formed chimeric spheroids by mixing two populations of Lung KP cells expressing a nuclear fluorescent protein (H2B-YFP) or mCherry-CAAX typically in a 1:10 ratio (Fig. 3C; Video S4). We can then easily track H2B-YFP-positive as they are much sparser, and their nuclear label is easy to detect. Using automated image analysis tools, we tracked hundreds of cells from different spheroids along the gradient of ischemia. Analysis of nearly 900 tracks of individual cells showed a large heterogeneity in their motility where some cells wandered near their origins, while others moved over large distances. On average, however, cells from ischemic clusters traveled further from their original location than cells from well-nurtured clusters (Fig. 3D, Fig. S3E,F; Video S4). This increase in net displacement was not due to an increase in speed (Fig. S3G) but rather because they moved more persistently than well-nurtured cells (Fig. 3E). Consistent with their increased persistence, the movement of ischemic cells was *superdiffusive* – *i.e.* significantly more directional than expected by a random walk^81^ (Fig. 3F, **m**ean **s**quare **d**isplacement (MSD) constant, σ=1.19; Fig. S3H). In contrast, MSD analysis of cells in well-nurtured clusters was indistinguishable from a random walk (Fig. 3F, MSD constant, σ=1.02; Fig. S3H).

**Figure 3:**
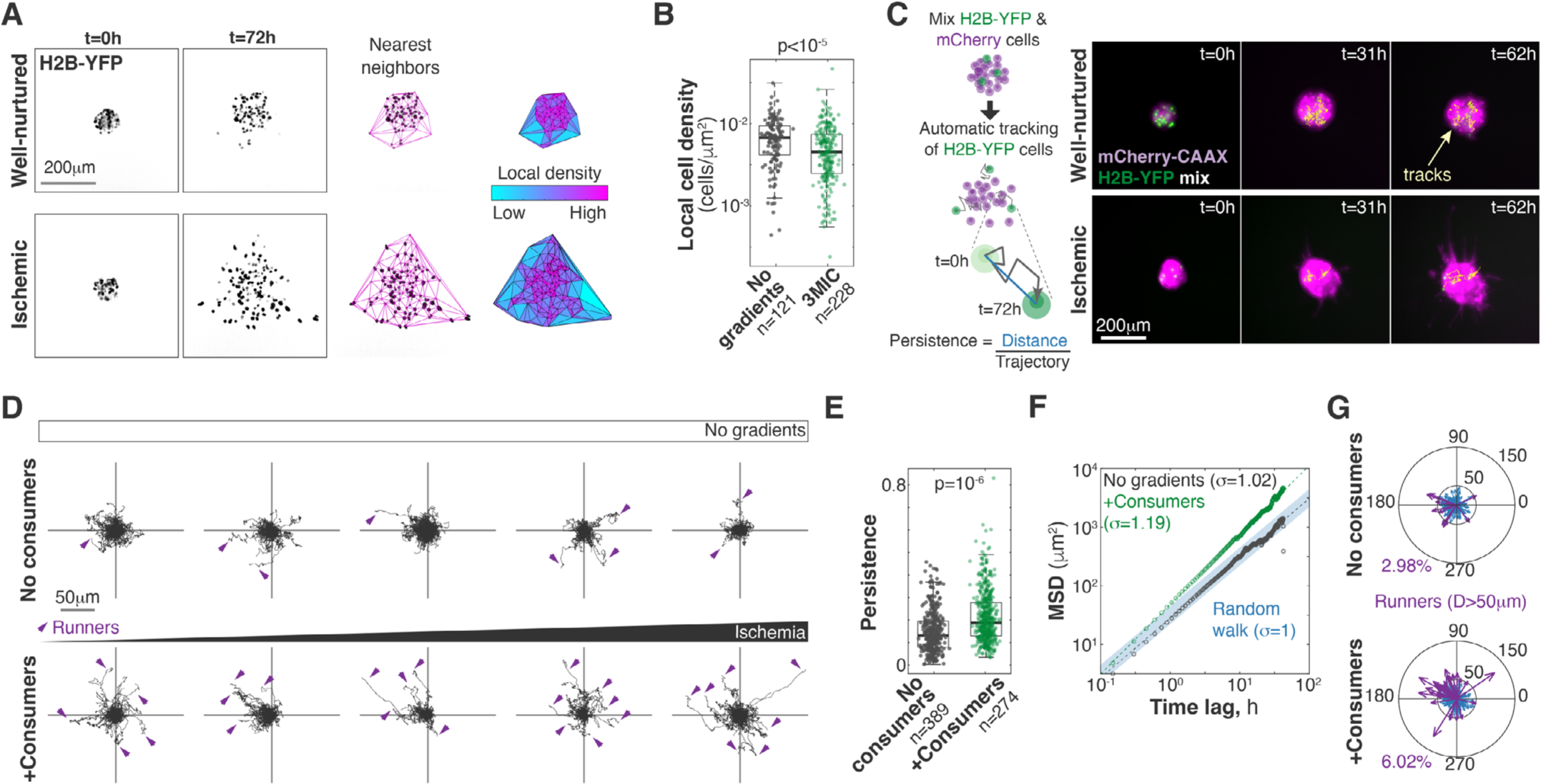
Ischemic cells in the 3MIC migrate more persistently. **A.** Image-based method to estimate local cell density. We located individual cells expressing the fluorescent nuclear marker H2B-YFP. We then triangulated closest neighbors and estimated areas between them. Local cell density corresponds to the inverse of each triangle’s area. **B.** Quantification of local cell densities over time for spheroids in the presence or absence of consumer cells. Data from 10 different spheroids (5 per condition). **C.** Experimental setup and representative data of automatically tracked individual cells. **D.** Top: migratory tracks of cells from 5 different tumor spheroids located at increasing distances from the opening of the 3MIC in the absence of consumer cells. Bottom: migratory tracks of cells from 5 different clusters located at similar distances from the opening as plots above but in the presence of consumer cells. Arrowheads show tracks for runner cells (defined as having a net displacement of 50*μ*m or more). **E.** Persistence of individual cells. Cells in ischemic spheroids are significantly more persistent than cells in well-nurtured spheroids. Data obtained from single cell tracks. **F.** Mean-squared displacement (MSD) analysis of all tracked cells. Ischemic cells disperse more than what would be expected from a random walk (MSD constant α>1). **G.** Compass plots showing the direction and magnitude of final displacement for individual cells. Runner cells are highlighted in purple. 3MIC opening (higher nutrients/oxygen levels) is to the left. Data from a representative experiment of three biological replicates.

To further explore these data, we defined *runners* as cells whose net displacement was greater than an arbitrary distance threshold (50µm). At most values of this threshold, ischemic clusters had about twice as many runners than well-nurtured spheroids or spheroids in chambers without consumer cells. Interestingly, we observed a directional bias in the movement of runner cells away from ischemia as if they were moving toward nutrient sources (Fig. 3G; Fig. S3I). However, this directional bias was small and only significant under some definitions (distance thresholds) of runner cells (Fig. S3J). We thus concluded that – at least in this setting – metabolite gradients do not act as significant orientation cues. Still, our data shows that ischemia leads to more persistent movements in three-dimensional tumor cultures resulting in highly dispersive and invasive tumor spheroids.

### Ischemic tumor cells lose epithelial features and increase ECM degradation

We then sought to determine specific cell changes triggered by ischemia. From a biophysical standpoint, the MSD changes observed in our cell tracking analyses, suggest that the movements of ischemic cells are less constrained^82^. Typically, constrains in cell migration are imposed by the ECM or by adhesion to other cells^42,43^. Consistently, hypoxic tumor cells are known to increase ECM invasion^23,83^ and to have lower levels of epithelial cell adhesion molecules such as E-Cadherin (E-Cad)^84–90^. We thus wanted to test whether ischemic spheroids increase ECM degradation and/or they show changes in cell-cell adhesion. To measure ECM degradation, we first used an assay where ECM proteins are laid over a coverslip coated with fluorescent gelatin^91^. More invasive clusters will break down the ECM network, and eventually degrade the labeled gelatin resulting in local loss of fluorescence (Fig. 4A). With this assay we observed that ischemia triggered a large increase in ECM degradation in tumor spheroids (Fig. 4B) and in consumer cells (Fig. S4A). As an orthogonal approach, we cultured tumor spheroids in DQ-Collagen – a fluorescent but quenched form of collagen that lights up when cleaved^92^. In this assay, ischemic spheroids also showed significantly higher levels of ECM degradation and foci of ECM degradation often colocalized with protrusive regions within spheroids (Fig. S4B).

**Figure 4:**
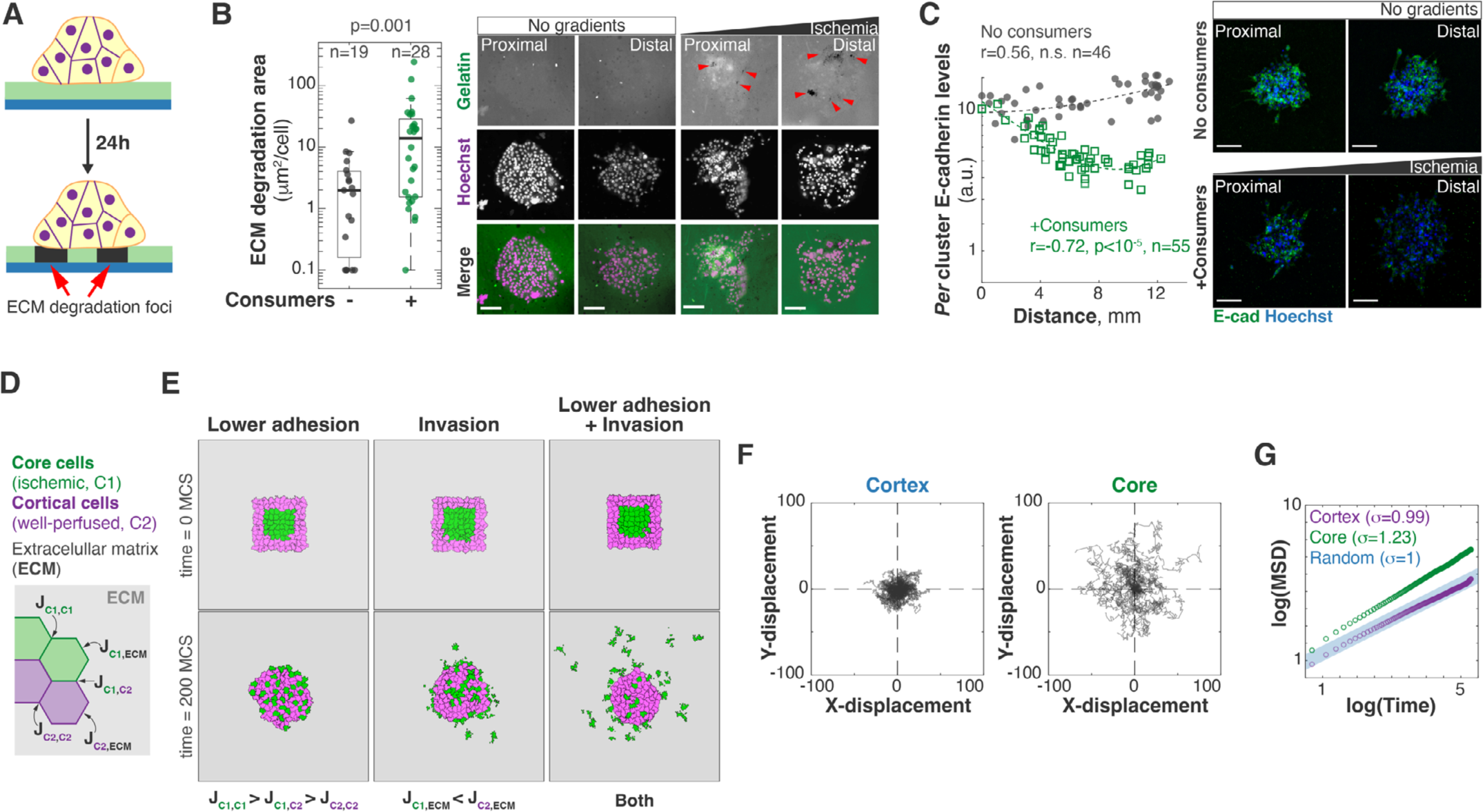
A decrease in epithelial adhesion and increased invasion can synergize to efficiently disperse tumor cells. **A.** Schematic depicting the ECM degradation assay where spheroids are grown on a glass coated with fluorescent gelatin. Local loss of fluorescence indicates ECM degradation foci. **B.** Representative images and quantification of ECM degradation. Spheroids were grown in the presence or absence of consumer cells. Ischemic spheroids show higher gelatin degradation (dark spots, red arrowheads). Bars: 100µm. Images compare spheroids located 2mm or closer to the opening of the 3MIC (Proximal) or further than 8mm (Distal). Data from a representative experiment of three biological replicates. **C.** Representative images and quantification of Lung KP spheroids grown in the 3MIC and stained for E-cadherin (E-cad, green). Nuclei are stained with Hoechst (blue). Bars: 100µm. Images compare spheroids located 2mm or closer to the opening of the 3MIC (Proximal) or further than 8mm (Distal). Data points: *per* spheroid E-Cad levels. Dashed line: linear fit. Data from a representative experiment of three biological replicates. **D.** Schematic depicting cell-cell and cell-substrate interaction energies from our Cellular Potts Model. See more details about the model on the main text and in the supplementary information. **E.** Initial and final conditions of simulated tumor populations moving under different rules. The combination of lower epithelial adhesion and invasion, but not each process alone, efficiently increase the dispersion of ischemic (core) cells. **F.** Tracks of simulated ischemic (core) and well-nurtured (cortical) cells modeled to have lower epithelial adhesion and increased invasion. **G.** MSD of simulated well-nurtured and ischemic cells with lower epithelial adhesion and invasion. MSD changes are in the same order of magnitude as in our experimental data (Fig. 3F).

Metabolic stress in the tumor microenvironment can decrease epithelial features such as E-Cad levels in carcinomas as we have previously shown in the MEMIC^71^. Consistent with these previous data, immunofluorescent analysis of ischemic spheroids in the 3MIC showed a dramatic decrease in E-Cad levels (Fig. 4C). We observed a similar trend using a fluorescent protein driven by the promoter of E-Cad (Fig. S4C), and thus suggesting that ischemia regulates E-cad at the transcriptional level. As many carcinoma cells, Lung KP show an intermediate EMT status mostly epithelial features including the expression of (Fig. S4D). While in our experiments we see a decrease in epithelial features, we did not observe significant changes in the levels of vimentin along the 3MIC suggesting that ischemia promotes a partial EMT (Fig. S4E). Interestingly, a recent study demonstrated that partial EMT in vimentin-positive carcinoma is required for metastasis in triple-negative breast cancer pre-clinical models^93^. In summary, these results show that ischemic cells decrease epithelial adhesion and increase the degradation of the surrounding ECM.

### A model of persistent tumor migration in the absence of directional cues

Our results are consistent with well-established evidence that tumor ischemia increase migratory and invasive cell properties^11,15,17,20^. Since these conditions arise deep within tumors, how do nascent metastatic cells move through the tumor tissue to eventually reach the tumor’s edge? An appealing hypothesis is that nutrients or oxygen act as directional cues that orient the movements of ischemic cells. However, we did not find enough evidence in our data to support this hypothesis (Fig. 3G; Fig. S3H,I). We thus asked whether our other observations: a decrease in epithelial adhesion and increased ECM degradation are sufficient to allow the escape of ischemic cells from the nutrient-deprived regions they emerge from.

We decided to address this question conceptually using a Cellular Potts Model^94,95^ – a mathematical framework frequently used to model problems of differential cell adhesion and cell sorting in morphological processes^96^. In a Cellular Potts Model, any number of ‘cell types’ exist on a grid and have different affinities for each other and for their surrounding substrate. High affinity contacts have a lower surface energy and thus they are more stable. In contrast, low cell-cell or cell-substrate affinities are represented as highly energetic and unstable contacts. At every simulated time point (a Monte Carlo Step, MCS), the model tries to minimize the energy of the system resulting in changes in cell shape, localization, etc. according to local energy levels^94,95^.

Our modeled tumors were formed by two cell types: *core* (ischemic) and *cortical* (well-perfused) cells designated according to their positions at the beginning of the simulation. Our specific goal was to test whether changes in cell-cell and cell-substrate adhesion would allow core cells to move through cortical cells and eventually invade into the surrounding ECM. We modeled the reduced levels of epithelial adhesion we observed in ischemic cells as a decreased cell-cell affinity by increasing in the surface energy between core cells i.e. decreasing their stability. Energy levels were set to ensure that the adhesion between core cells was low, between cortical cells was high, and intermediate between the two cell types. We tested a range of energy parameters obtaining similar qualitative results as long as we kept these relationships (Fig. 4D; see additional details in Tables S1 and S2). Implementation of these changes resulted in clusters where core cells mixed with cortical cells but there was no invasion into their surroundings (Fig. 4E; Video S5). To model the increased ECM invasion in ischemic cells, we increased the affinity for the surrounding ECM in core. The intuition behind this step is an increase in ECM degradation will facilitate the movement of tumor cells into the ECM, which in a Cellular Potts Model equates to a higher ECM-tumor affinity (Fig. S4F). This feature alone, produced poor mixing between populations and a moderate trend to invade the ECM (Fig. 4E; Video S5). However, cell movements were random and did not show the persistence we observed in our experimental data (Fig. S4G).

The combination of these two rules however – core cells with decreased cell-cell adhesion and increased affinity for the ECM – produced the most interesting outcomes. Ischemic core cells moved through layers of well-perfused cells and rapidly dispersed through the surrounding ECM (Fig. 4E; Video S5). These virtual cell movements were remarkably close to the movements of our experimental cells. For example, tracks of core and cortical cells in the model (Fig. 4F), resembled the movements of ischemic and well-nurtured cells, respectively (compare with Fig. S3F). Most strikingly, the MSD of core cells in this scenario increased by the same order of magnitude as in our experimental cells in the 3MIC (compare Fig. 4G and Fig. 3F). Altogether these data suggest that increasing the affinity for the ECM and decreasing cell-cell adhesion is sufficient for the spontaneous emergence of directional movements driving the dispersion of core cells into the ECM – even in the absence of explicit directional cues. While many other mechanisms will contribute to the translocation of ischemic tumor cells to regions from where they can spread from *in vivo*, this model proposes a minimal set of conditions that can facilitate this process.

### Pro-metastatic features induced by ischemia are reversible

Hypoxic and ischemic environments can select for more aggressive and metastatic clones^54,97,98^. The contribution of clonal selection in our brief 3MIC experiments (∼24-72h) is negligible. *In vivo* however, time scales are much longer, which may allow ischemia to select for more migratory clones. We hypothesized that if ischemic cells are selected to be more migratory *in vivo*, they should display a higher migratory phenotype in the 3MICs – irrespective of their location along the ischemic gradient. We thus injected Lung KP cells expressing HRE-GFP into the flank of syngeneic mice. After 2 weeks, we extracted these tumors and dissociated them into single cells. Using flow cytometry, we sorted ischemic from non-ischemic cells according to their GFP levels (Fig. S5A). From these sorted cells, we generated tumor spheroids, which we then cultured into separated 3MICs (Fig. 5A; Fig. S5B). As shown in figure 5, the increase in cell invasion strongly correlated with the distance to the opening but not with their precedence (Fig. 5B,C). Consistently, spheroids formed neither from hypoxic nor from non-hypoxic tumor cells showed and increased invasion when cultured in 3MICs without consumers (Fig. S5C,D). These results show that, at least in this system, the pro-metastatic features produced by ischemia are phenotypic changes that do not require clonal selection.

**Figure 5:**
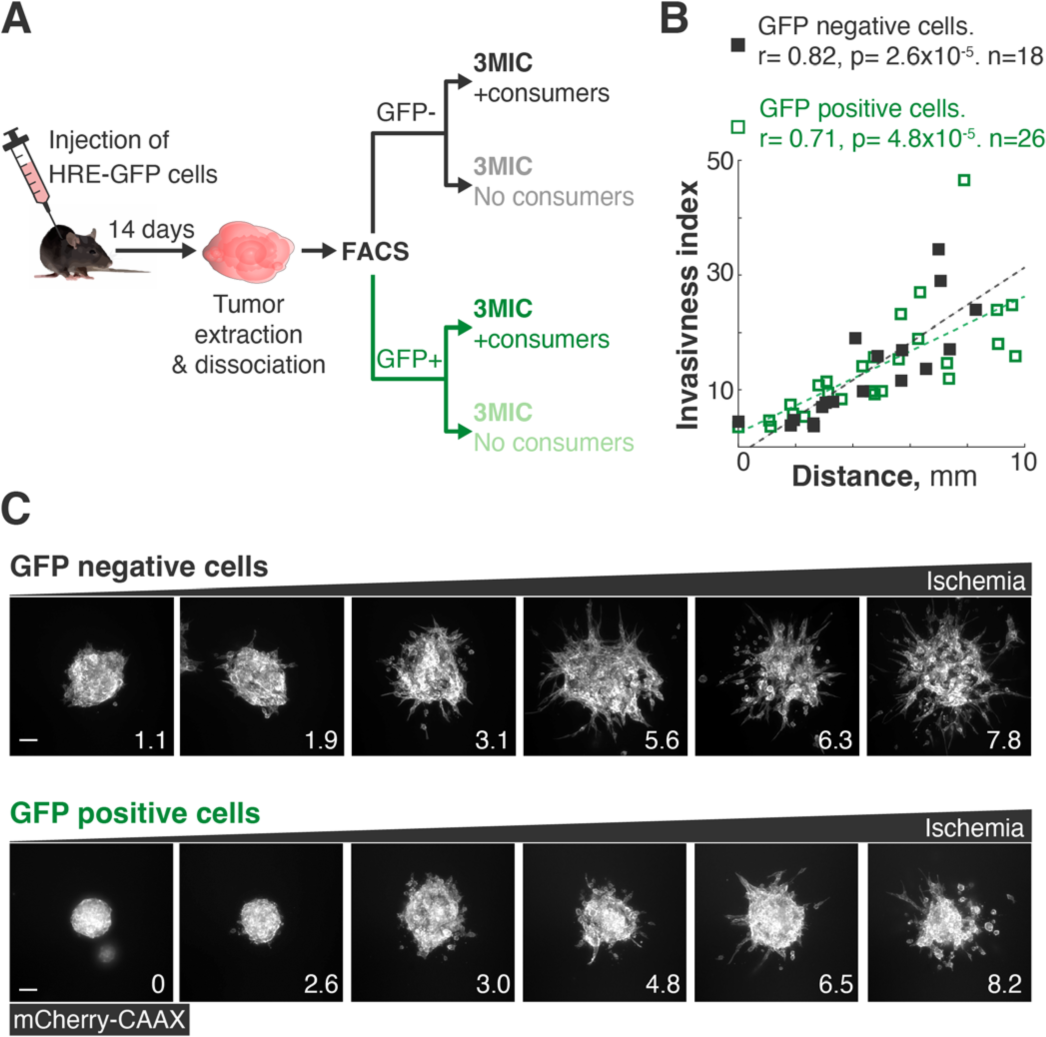
Tumor hypoxia *in vivo* did not induce permanent changes in the migratory potential of tumor cells. **A.** Lung KP cells expressing HRE-GFP and mCherry-CAAX were injected subcutaneously in C57BL/6 mice and allowed to grow for 14 days. Tumors were then extracted and sorted into GFP positive (hypoxic/ischemic) and GFP negative (normoxic/well-nurtured) populations. These different sub-populations were grown as tumor spheroids in 3MICs in the absence or presence of consumer cells. **B.** Quantification of invasiveness of experimental setup shown in (**A**). Data points: invasiveness index of individual spheroids. Dashed lines: linear fit. We observed similar invasion levels and response to ischemia in all spheroids regardless of if they were derived from well-nurtured or ischemic tumor regions. **C.** Representative images of tumor spheroids derived from GFP positive and GFP negative cells growing at different locations inside of 3MICs. Numbers in lower right corner denote the distance to the opening in mm. Bars: 100*μ*m.

### Reconstruction and visualization of more complex microenvironments

Solid tumors are infiltrated by a variety of stromal and immune cells^22,29,30^. Tumor-associated macrophages (TAMs) are one of the most abundant cell types in solid tumors and their presence correlates with increased cancer progression, metastasis, and mortality^20,30,31,99–102^. TAMs recruit endothelial cells helping to orchestrate the vascularization process required for tumor growth and metastasis^31^. We recently showed that a combination of hypoxia and high lactic acid, activates the ERK/MAPK pathway in macrophages who then secrete VEGFA and induce tube-like morphogenesis in endothelial cells^70^. To test whether we could recapitulate these data in the 3MIC, we generated endothelial clusters from SVEC4-10 cells (referred as SVECs) and co-cultured them with bone marrow-derived macrophages (BMDMs). As controls we examined similar SVEC clusters in 3MICs without macrophages. In the absence of macrophages, clusters of endothelial cells remained mostly rounded, even in the presence of consumer cells and regardless of their location along the ischemic gradient (Fig. 6A). In the presence of macrophages however, ischemic endothelial cells extended away from clusters and sprouted into the ECM (Fig. 6A; Video S6). This tube-like morphogenesis did not occur in well-nurtured regions of the 3MIC (Video S6). We did not notice that the cell type used as consumers produced significant differences in endothelial sprouting, but it always required the infiltration of macrophages (Fig. S6A). Consistent with previous evidence^31,70^, inhibition of VEGF signaling with linifanib abrogated the endothelial sprouting induced by macrophages (Fig. 6B; Fig. S6B). While this inhibitor may have off target effects, these data are consistent with prior data showing VEGF secretion and a pro-angiogenic role ischemic in ischemic macrophages^31,70^.

**Figure 6:**
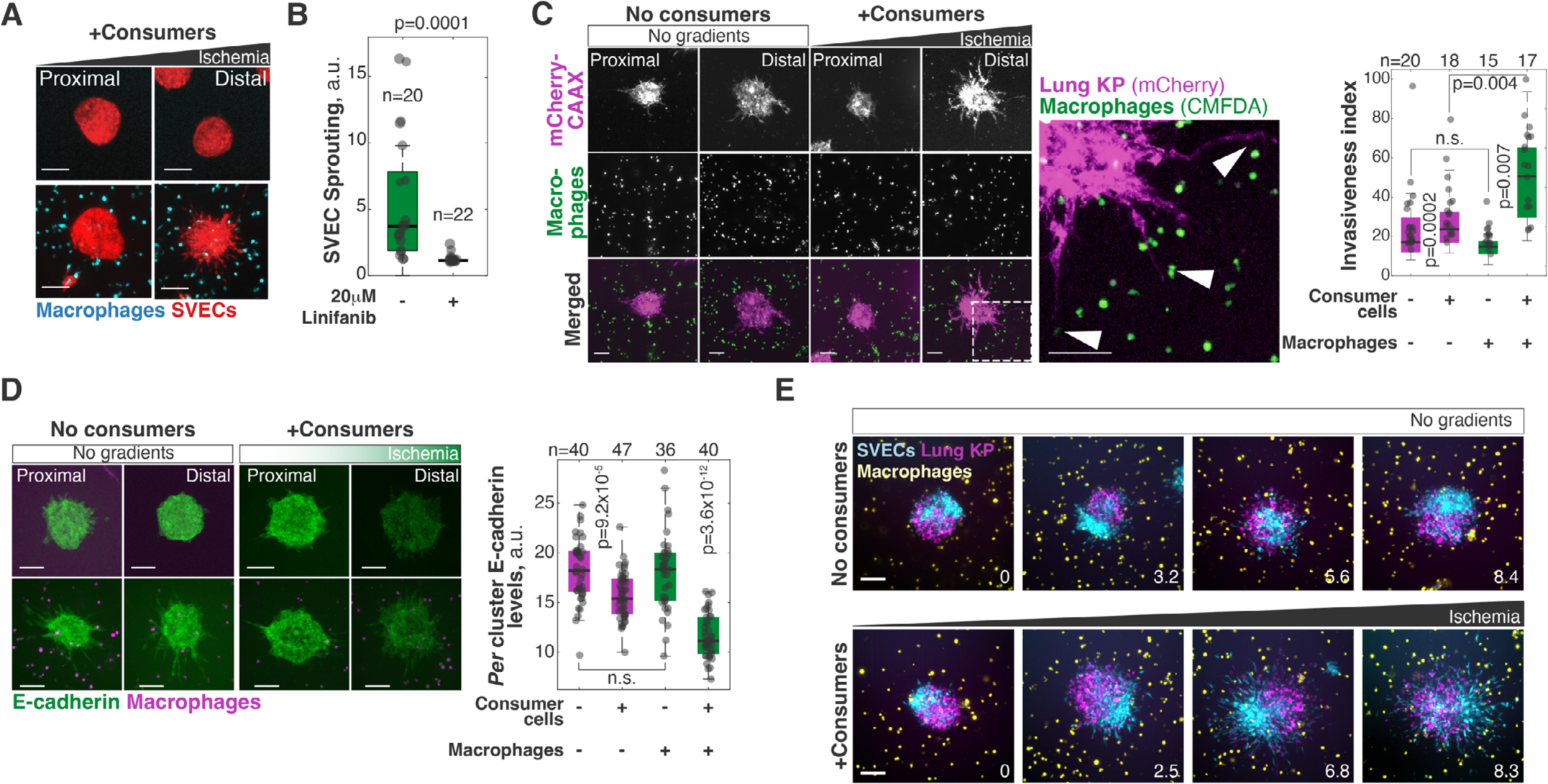
The 3MIC allowed the seamless implementation of cell co-cultures. **A.** Clusters of endothelial cells (SVECs, red) grown in 3MICs in the presence or absence of bone marrow-derived macrophages and stained with CMFDA (cyan). Bars: 100µm. Images compare spheroids located 2mm or closer to the opening of the 3MIC (Proximal) or further than 8mm (Distal). Data from a representative experiment of four biological replicates. **B.** Quantification of SVEC sprouting in clusters co-cultured with macrophages. VEGFA inhibitor Linifanib abrogates sprouting induced by ischemic macrophages. Data from a representative experiment of two biological replicates. **C.** Representative images and quantification of invasiveness of Lung KP spheroids co-cultured with BMDMs in the 3MIC. Ischemic macrophages significantly enhance tumor invasiveness and synergizes with ischemia. Region delineated with dashed lines is magnified on the right. Data points: *per* spheroid invasiveness. Bars: 100µm. Images compare spheroids located 2mm or closer to the opening of the 3MIC (Proximal) or further than 8mm (Distal). Data from a representative experiment of three biological replicates. **D.** Representative images and quantification of Lung KP spheroids grown in the 3MIC and stained for E-cadherin (E-cad, green) in the presence or absence of macrophages (magenta). Ischemia and the presence of macrophages synergize in their reduction of E-Cad levels. Bars: 100 µm. Images compare spheroids located 2mm or closer to the opening of the 3MIC (Proximal) or further than 8mm (Distal). Data from a representative experiment of four biological replicates. **E.** Representative images of triple co-cultures: chimeric Lung KP cells and SVEC spheroids, were grown in the presence of macrophages with or without consumer cells. Numbers in lower right corner denote the distance to the opening in mm. Bars: 100*μ*m.

We then co-cultured macrophages with tumor spheroids. In these experiments, we observed that the presence of macrophages increased the migration and ECM invasion of tumor cells. This increase was evident even in well-nurtured regions, but much stronger under ischemia (Fig. 6C). Similarly, macrophages and ischemia synergized to decrease E-Cad levels (Fig. 6D).

Live imaging of these tumor-macrophage interactions also revealed a fascinating behavior: tumor cells from ischemic spheroids extended protrusions towards macrophages appearing to physically drag them into the tumor cell cluster (Fig. 6C; Video S7). This observation suggests that there may be an understudied mechanical interaction where malignant cells may ‘pull’ macrophages into the tumor. Increasing the complexity of co-cultures in the 3MIC is simple. For example, we produced a triple co-culture of chimeric SVEC/Lung KP spheroids grown in ECM and surrounded by macrophages (Fig. 6E; Video S8). Here again we observed the strong synergy between metabolic and macrophage signals in promoting morphological changes in tumor and endothelial cells.

### Using the 3MIC to test the effects of anti-motility drugs on ischemic cells

Limited diffusion of anticancer drugs from blood vessels into solid tumors is a major therapeutic challenge^103,104^ (Fig. 7A). Multiple factors account for poor drug penetration including tumor vascularization and the distance between blood vessels^105^, simple diffusion through of cells and the ECM^106^, and drug consumption and degradation by cells in the tumor microenvironment^107^. In addition, hypoxic and ischemic cancer cells can develop drug resistance by increasing ABC transporters and drug efflux^108^, and bypass drug targets through metabolic rewiring^109^. Distinguishing biophysical limitations to drug penetration from the acquisition of true drug resistance is necessary to improve therapy and yet most assays struggle to untangle these factors^104^. The 3MIC can solve some of these issues providing a unique opportunity to test drugs that directly target metastatic initiation.

**Figure 7:**
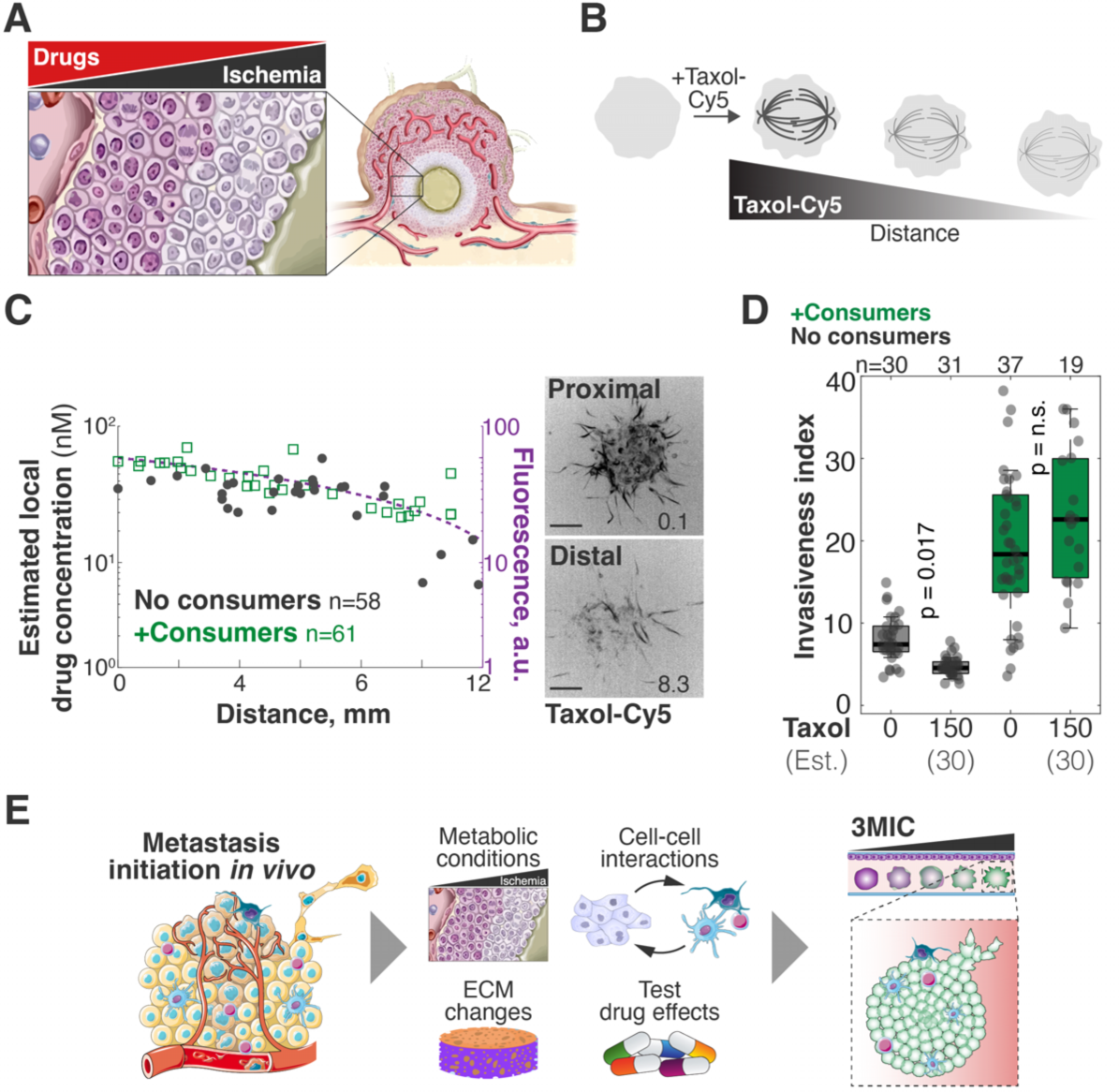
The 3MIC allows studying the effects of anti-migratory drugs on ischemic cells. **A.** Cartoon showing that drug penetration of tumors is a therapeutic challenge, especially in ischemic regions where drug levels are likely lower. **B.** Taxol-Cy5 is a probe used to fluorescently label microtubules. We used this property to estimate how Taxol levels decrease due to diffusion along the 3MIC. **C.** Quantification of Taxol-Cy5 in spheroids within 3MICs with or without consumer cells. Plots indicate that drug levels decrease as they diffuse through 3MICs regardless of the presence of consumer cells or not. Dots: Taxol-Cy5 and location of single spheroids. Data pooled from two biological replicates. **D.** Invasion of Taxol-treated spheroids growing in 3MICs with or without consumer cells. Cultures were treated with 150nM of Taxol which we estimated to drop to about 30nM deeper within 3MICs. Points: invasiveness of individual distal clusters (located at 4mm or more from the opening). Data from a representative experiment of three biological replicates. **E.** Cartoon showing features of the 3MIC as an *ex vivo* model to study metastasis.

To illustrate this principle, we treated tumor spheroids with anti-migratory doses of Taxol. At high doses, Taxol stabilizes microtubules leading to mitotic arrest and cell death^110^. However, sub-lethal concentrations of this drug inhibit migration and invasiveness in several cell types^111,112^ (Fig. S7A). In our experiments, we determined that concentrations of 20-200nM of Taxol drastically inhibited the movements and invasion of well-nurtured KP lung spheroids with no effects on their cell viability (Fig. S7B). Similar levels of this drug however, had much milder effects or no effects at all on the movements of ischemic tumor spheroids (Fig. S7C).

To distinguish whether these differences are due to a decrease in Taxol levels along the 3MIC, or if they denote a differential response in ischemic cells, we took advantage of fluorophore-conjugated Taxol analogues. These compounds are routinely used to label microtubules, and thus we can use the intensity of microtubule staining as a proxy for local drug levels (Fig. 7B; Fig. S7D). After adding known quantities of the fluorescent Taxol analog (Taxol-Cy5), fluorescence levels progressively decreased in distal spheroids, but they did so at the same rate in 3MICs with or without consumer cells (Fig. 7C; Fig. S7D,E). These results indicate that consumer cells have a negligible effect on the Taxol gradient. From the changes in microtube staining intensity, we determined that in a 3MIC culture treated with 150nM of Taxol, the most ischemic spheroids will experience about ∼30nM of Taxol (45nM at 10mm and 24nM at 12mm, Fig. 7C). While these Taxol levels have no effects on ischemic tumor spheroids, 30nM of this drug completely inhibited the movements of well-nurtured tumor cells (Fig. 7D). We thus concluded that ischemic spheroids are more resistant to Taxol by mechanisms beyond drug penetration and diffusion.

## DISCUSSION

Animal models are fundamental to study the complexity and heterogeneity of the tumor microenvironment^113,114^. In these *in vivo* models however, it is hard to isolate and weigh the contributions of different variables to tumor progression, and large experiments are prohibitively expensive for most laboratories^61,62^. On the other hand, conventional *in vitro* approaches offer much better experimental control and can be easily scaled up for high throughput approaches. However, *in vitro* models do not usually consider the cellular and molecular heterogeneity of the tumor microenvironment. The lack of models that can bridge the complexity of *in vivo* models with the ease of *in vitro* cultures has impaired progress in understanding the tumor microenvironment and how it can modulate the emergence of metastatic tumor features^2,115,116^. To bridge this technical gap, we described the design and implementation of the 3MIC – an *ex vivo* culture system that allows visualizing the acquisition of metastatic-like migratory properties in complex 3D tumor cultures. Altogether the experiments presented here show that the 3MIC recapitulates key features of the tumor microenvironment. It allows for imaging ischemic cells that are hidden to conventional techniques and permits testing the role of different elements of the tumor microenvironment and the effects of drugs on the emergence of metastatic features (Fig. 7E).

The results shown here are consistent with well-established evidence that hypoxia is essential for the emergence of metastasis. The 3MIC however, allowed us to gain new insights into this process by 1) re-creating a complex ischemic-like metabolic microenvironment rather than simple hypoxia and 2) overcoming challenges associated with imaging ischemic cells that are usually located deep within tumor structures. The ischemic-like environment formed in the 3MIC better mimics metabolic conditions in tumors such as accumulation of lactic acid and nutrient deprivation^52,71,117^. In our experiments, we found evidence that media acidification has a more direct effect on the invasive properties of ischemic spheroids than hypoxia. We think that media acidification may be the direct signal that triggers invasion under ischemia while hypoxia indirectly contributes to this effect through the pro-glycolytic role of HIF1A^78^. At this moment, we can only speculate why pH perturbations may have such strong effects in our experiments. A possible explanation is that many ECM-digesting metalloproteases have lysosomal origins or are more active in more acidic environments and thus extracellular acidification would increase the efficiency of these enzymes^43,77,118^. While our data shows a critical role for hypoxia and extracellular pH, additional metabolic^37,38^ and non-metabolic cues are likely to increase cell invasion under ischemia. For example, glucose deprivation triggers the release of pro-migratory cytokines^119^ and there is a growing list of metabolites that directly or indirectly affect the number and destination of metastases^37,38^.

Our data shows that ischemic cells display a more efficient dispersion. Consistent with previous evidence^120,121^, we showed here that ischemia decreases epithelial cell adhesion and increases the degradation of the extracellular matrix. While some of our observations share features with the process of epithelial-to-mesenchymal transition (EMT), we preferred referring to a partial EMT as we only observed a decrease in epithelial features without a significant change in mesenchymal markers. In addition, the contribution of EMT to the metastatic process remains controversial^86,122,123^. An additional issue to consider is the composition of the ECM as they can alter the phenotypes and behavior of tumor and stromal cells^124^. While in our experiments we used a simple combination of collagen and Matrigel in our experiments, the 3MIC allows testing different ECM compositions and structural scaffolds (Fig. 7E).

At the single-cell level, ischemia increases cell persistence without a significant change in cell polarity or increasing directional cell migration. However, we observed a small bias towards higher nutrients in the direction of cell movements. While in our setting these differences were not statistically significant, this observation would be consistent with a widespread set of examples of nutrient-driven chemotaxis in prokaryotes and in other eukaryotic cells^125,126^. For example, the key nutrient sensor mTORC2 is critical in neutrophil chemotaxis. Still, additional experiments will be needed to establish whether or not tumor cells can follow nutrient gradients.

Recent evidence shows that tumor hypoxia can act as an evolutionary pressure that selects for tumor clones that are more resilient to stressors such as reactive oxygen species^86,97,127^. However, our data shows that hypoxia and ischemia can alter cells directly without the need for clonal selection. These two ideas are not in conflict and our experimental conditions are not free of caveats. Most likely the acquisition of metastatic features in patients occurs by a combination of selection and opportunistic adaptations to the tumor microenvironment.

The direct visualization of tumor-stroma interactions afforded by the 3MIC has unique advantages. For example, we were excited to observe that the movement of macrophages toward tumor structures seemed to be driven by tumor cells mechanically dragging macrophages into their clusters (see for example Video S7). It is possible then, that immune infiltration of tumors is aided by physical recruitment by cancer cells. While we are not aware of previous reports of this behavior, this may be due to the challenges of observing tumor-macrophage interactions *in vivo*, especially in ischemic tumor regions.

It has been difficult to discern whether a decrease in drug response *in vivo* is due to lower drug concentration or to intrinsic changes that make cells more drug-resistant^128^. This is particularly true in more fibrotic and ischemic tumors such as pancreatic ductal adenocarcinomas^129,130^. The 3MIC however allows for the separation of these effects and we were able to show that ischemic cells display a true Taxol resistance mechanism. These experiments illustrate the ease of testing drugs on ischemic cells and untangling biophysical factors such as drug diffusion from biological adaptations to anticancer drugs. While the particular example we used benefited from the availability of labeled Taxol analogs, a similar approach is possible using imaging mass spectrometry and other methods that allow the quantification of local drug levels^131^.

Finally, the fabrication of the 3MIC is easy and affordable with a conventional 3D printer. This system can seamlessly integrate existing co-culture and *ex vivo* protocols and thus, we invite the research community to use the 3MIC when studying metastasis and other processes where the limitation of nutrients in 3D multicellular structures is relevant.

## METHODS

### Cell culture

Lung KP clones and Lung KPKeap1 clones were derived from the KRAS^G12D^/TP53^-/-^ and KRAS^G12D^/TP53^-/-^/Keap1^-/-^lung adenocarcinoma model, respectively, developed and kindly shared by Dr. Thales Papagiannakopoulos. MCF10A, MCF7, DlD1, and SVEC-4-10 cells were purchased from ATCC. C6-HRE-GFP cells were a kind gift from Dr Inna Serganova (Memorial Sloan Kettering Cancer Center, New York, NY, USA). Cells were cultured in High Glucose DMEM (Gibco, #11965-092) supplemented with 10% Fetal Bovine Serum (FBS; Sigma-Aldrich, #F0926) at 5% CO_2_ and 37°C. MCF10A cells were cultured in High Glucose DMEM (Gibco, #11965-092) supplemented with 5% Horse Serum, 20ng/ml Animal-Free Recombinant Human Epidermal Growth Factor (Peprotech,#AF-100-15), 0.5 mg/ml Hydrocortisone (Sigma-Aldrich, #H0888), 100 ng/ml Cholera Toxin (Sigma-Aldrich, C8052), 10μg/ml Insulin, at 5% CO_2_ and 37°C. To induce HIF1A signaling in normal oxygen conditions, cell spheroids were treated with 2mM DMOG (Selleckchem, #S7483) or with 300µM CoCl_2_ (ThermoFisher Scientific #012303-30) for 18 hours. To alter the pH of the media, we added 25mM of NaHCO₃ (ThermoFisher Scientific #25080094) or 20mM of lactic acid (Sigma #L1750) to the media.

Lung KP cells stably expressing a hypoxia reporter (pLenti-5XHRE-GFP, Addgene #128958, denoted here as HRE-GFP), cell membrane marker (pLenti-mCherry-CAAX, Addgene #129285), E-cadherin reporter (pHAGE-E-cadherin-RFP, Addgene #79603), cell cycle reporter (pBOB-EF1-FastFUCCI-Puro, Addgene #86849), LV-YFP (Addgene #26000), and pLenti-H2B-iRFP720 (Addgene #128961) were generated using standard lentivirus mediated stable cell line preparation protocols. H2B-mCherry was cloned by removing YFP from the LV-YFP vector using BamHI and KpnI restriction enzymes. The linearized vector containing H2B was then ligated with mCherry using SLIC^132^.

### Formation of 3D tumor spheroids

Cell spheroids were formed via hanging drops following standard protocols^133^. Briefly, cells were dissociated into a single-cell suspension of 10^4^ cells/ml. The cell suspension was distributed into 20µl droplets onto the lid of a Petri dish. The base of the dish was filled with PBS (Gibco, #14040-133) or distilled water to prevent droplet evaporation during the incubation time. The lid was then inverted onto the dish and incubated for 96 hours, to ensure the formation of compact spheroids. Typically, we cultured ∼50 spheroids in each 3MIC chamber.

### Isolation and differentiation of BMDMs

Bone marrow-derived macrophages (BMDMs) were extracted from C57BL/6 mice following standard protocols^69,70^. Following isolation of the bone marrow, cells were cultured in low attachment culture dishes (VWR, #25384-342) in High Glucose DMEM supplemented with 10% FBS and 10ng/mL Recombinant Mouse CSF-1 (R&D Systems, #416-ML) for 7 days.

### Establishment of tumor-derived clones

Approximately 5x10^4^ cells Lung KP cells, stably expressing pLenti-5XHRE-GFP and pLenti-mCherry-CAAX were injected subcutaneously into C57BL/6 mice and grown for 14 days. Tumors were then extracted and dissociated using a solution containing of 2U/ml Dispase and 4mg/ml Collagenase IV. Tumor cells were enriched as a CD45 negative population and sorted to their GFP levels. GFP-positive and negative cells were grown separately as clusters using the hanging drop method described above.

### Generation of Hif1a KO cells

Hif1a knockout cells were generated using CRISPR/Cas9 genome editing. Briefly, forward and reverse sequences (CACCGAGATGTGAGCTCACATTGTG and AAACCACAATGTGAGCTCACATCTC) were synthesized to form single guide RNAs (sgRNAs) targeting the Hif1a gene. A vector (lentiCRISPR-v2-puromycin, Addgene #98290) carrying this sgRNA was introduced into HEK293T cells together with envelope and packaging plasmids (VSV-G and Delta-8.9) via transfection using Lipofectamine (Thermo Scientific #L3000008). Target cells were then infected with collected lentiviral particles, selected using puromycin (10µg/mL Gibco #A1113803) and separated into single clones through limiting dilution.

### 3D printing and microfabrication

To create the framework of 3MIC fused filament fabrication was used. Designing of the framework was done using openSCAD (https://openscad.org). 3MICs were printed in two parts that were later assembled: the main framework and the upper coverslip holder (where consumer cells are grown). These parts were printed in an Ultimaker 3B printer using Black PLA filament. Alternatively, the framework can be printed using Stereolithography (SLA) printers. In this case, the entire framework (the upper coverslip holder and main framework) can be printed as one unit. We used Dental SG resin to print the framework on Form 3 (Formlabs). The uncured resin was removed from the prints by washing them in isopropanol for 1 h and further post-curing it with UV irradiation heated at 60°C. A glass coverslip (VWR, No. 1, 22x60mm, #48393-070) was glued to the base of the main framework using a UV-curable adhesive (Norland Products, NOA68) using a long-wave UV lamp (MelodySusie, DR-5401). Then, the coverslip holder fragment was glued onto the glass coverslip. Finally, the assembled 3MIC was sterilized in a short-wave UV lamp chamber (Meishida, CM-2009) for 15 minutes.

### Culture of tumor spheroids in the 3MIC

To prepare the layer of consumer cells, 24hrs before the day of the experiment, a coverslip (VWR #48366-045) was coated with Poly-D-lysine (Sigma-Aldrich, #P6407) to aid adherence of cells to the glass surface. Cells were detached using trypsin and diluted into a cell suspension of a density of 3.5x10^5^ cells/mL for faster dividing cells and 10^6^ cells/mL for slower dividing cells. The coverslip was placed into a well of a 6-well dish and covered with 1mL of the cell suspension. The coverslip was incubated overnight at 37°C to let the cells adhere to the coverslip. On the day of the experiment, the spheroids were collected from the Petri dishes by inverting the lid and gently flushing all the spheroids with DMEM supplemented with 10% FBS using a 5mL serological pipette. The spheroid containing suspension was collected and centrifuged at 50G for 10-15 minutes. Meanwhile, a thin bed of 110µL ECM was made in the 3MIC – here, we used either 2.2mg/mL rat tail Collagen (Corning, #354236) – or 2:3 mix of Matrigel and 1.6mg/ml of rat tail Collagen I and incubated at 37°C for 20 mins to allow the polymerization of the ECM. Spheroids were resuspended in about 110µL of ECM and immediately transferred onto the polymerized ECM bed. The ECM with spheroids was polymerized at 37°C for ∼30 minutes. With the help of forceps, the coverslip containing the consumer cells was inverted and inserted into the slot of the coverslip holder. The 3MIC was then gently filled with 1.25mL of media, avoiding trapping bubbles underneath the consumer coverslip. Control wells are assembled in the same way, but the top coverslip has no consumer cells.

### Collagen degradation assay

To investigate the invasive potential of cells across the gradient, the ECM was mixed with DQ-collagen, type I From Bovine Skin, Fluorescein Conjugate (Invitrogen, #D12060) at a final concentration of 25µg/ml. This ECM was used to prepare the bed and embed the spheroids as well. The extent of degradation of the matrix was determined by measuring the fluorescence intensity of cleaved collagen.

### Gelatin degradation assay

Glass coverslips were coated with gelatin from pig skin, Oregon Green 488 Conjugate (Thermo Fisher Scientific, #G13186), by inverting coverslips on 20μl drops of fluorescent gelatin (1mg/ml) and incubating for 20 minutes. The coverslips were then lifted, excess liquid drained, and inverted on 40μl drops of 0.5% glutaraldehyde for fixation. The coverslips were then washed with PBS and glued onto the PLA frame using the UV-curable adhesive (Norland Products, #NOA68) and cured in a long-wave UV lamp (MelodySusie, #DR-5401) for 15 minutes. The cured coverslip was then UV-sterilized and used for the experiment. Briefly, the spheroids were collected from the hanging drops and centrifuged and resuspended in 100μl of media. The spheroid solution was carefully placed on the coated coverslips and incubated for 30-60 minutes to allow the spheroids to settle down. After ensuring that there are no floating spheroids, the consumer coverslip was transferred into the 3MIC and the remaining 1.4ml of media was added. The 3MICs were incubated at 37°C, for 24 hours. After 24 hours, the 3MICs were fixed using 4% paraformaldehyde (PFA) for 10 mins and imaged using 10X objective of Nikon Eclipse Ti2-E spinning disk confocal microscope.

### Microscopy

Microscopic images were captured using a 10x objective of Nikon Eclipse Ti2-E spinning disk confocal microscope, at fixed timepoints or set up for time-lapse microscopy. For time-lapse microscopy, cells were maintained at 37°C using a stage top incubator (Tokai Hit, #INUCG2-KRi) and the chamber was connected to a humidified mixed gas source (5% CO_2_). Spheroids from different regions of the 3MIC were imaged at suitable time-intervals in different fluorescent channels. Typically, we imaged ∼10 spheroids per 3MIC every 15-30 mins for 24h. Extracellular pH was estimated using the ratio of the green over the blue channels one RGB images of cell culture media containing the pH indicator phenol red. As pH decreases and the media yellows, the signal on the green channel increases while the blue channel remains largely unchanged. To match this ratio to specific pH values, we imaged and measured the pH of the same culture media supplemented with different levels of lactic acid.

### Immunofluorescence

Spheroids within the 3MIC were fixed using 4% PFA (Affymetrix, #19943) for 10 minutes at room temperature and washed four times with PBS. Cultures were permeabilized using PBS containing 0.5% (vol/vol) Triton-X-100 (Sigma-Aldrich, #T8787) for 20 minutes and washed three times with PBS. Slides were blocked in 10% FBS in immunofluorescence (IF) buffer (0.3M glycine in PBS) for 60 minutes at room temperature. Primary antibodies were diluted at 1:200 in the same blocking solution. We used anti-phospho-H3 (Cell Signaling Technologies, #9706), E-cadherin (BD Bio Sciences, #610182), and CD45 (R&D, #AF114) antibodies. 3MICs were incubated with primary antibodies overnight at 4°C, then washed with IF buffer and incubated overnight at 4°C with secondary antibodies also diluted in blocking solution at 1:200. Cultures were then washed and counterstained with 0.5μg/mL Hoechst 33342 in PBS (Invitrogen, #H1399).

### Image analysis

Automated cell tracking was conducted as previously described^69^. Persistence is defined as the ratio between the net displacement of a cell over its trajectory. We estimated the local cell density using a method based on the Delaunay triangulation of neighboring cells^69,70^. Per cell, per spheroid, and population levels of florescence were estimated using custom image analysis software as previously described^71^.

### Cellular Potts Model (CPM) simulations

Simulations were conducted using a MATLAB CPM implementation^134^. Adhesion between core (C1) and cortical (C2) cells were implemented as:

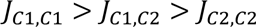

Where *J_i,j_* is the surface energy at every contact point between the two cells^95^. This relationship modeled the decrease in E-Cad levels in ischemic cells. The differential affinity for the ECM in core cells was simulates as:

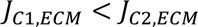

The CPM was implemented in a 200x200 grid with 100 initial cells and run for 1,28x10^8^ Monte Carlo Simulation steps. Parameters used in the conditions presented here can be found in Tables S1 and S2. MATLAB routines to run the simulations are available upon request.

### Statistics

We conducted at least three biological replicates of all experiments, unless stated otherwise. Unless noted, we used two-tailed t-student tests to estimate p-values between two conditions and Pearson’s linear coefficient when testing correlations. For multiple comparisons, we used one-way ANOVA followed by a Tukey-Kramer test. In all plots, error bars show standard deviation from the mean. In boxplots, center lines show the median and box edges show 75^th^ and 25^th^ empirical quartiles of the data.

## ACKNOWLEDGEMENTS

We thank all members of the Carmofon Laboratory for critical discussions about this project and the manuscript. This work was supported by awards to CC-F from the National Cancer Institute of NIH (DP2 CA250005) and the American Cancer Society (RSG-21-179-01-TBE). CC-F is a Pew Scholar in the Biomedical Sciences, supported by The Pew Charitable Trust (00034121). JG was supported by a NIH Training Grant (1T32GM132037-01).

## Authors contributions

Concept and experimental design CC-F with help from LA. Experiments by LA, JG, JL, MR, and C-CF. Image analysis: CC-F, LA, and LJ. Mathematical modeling: CC-F. Manuscript written by CC-F with feedback and contributions from all authors.

## Data and materials availability

All data is available in the manuscript or the supplementary materials. Original raw files and analysis software are available upon request. Use of the 3MIC requires a Material Transfer Agreement and it is free for non-commercial and academic uses.

## ADDITIONAL INFORMATION

Supplementary information is available for this paper. Correspondence and requests for materials should be addressed to CC-F.

## Supplementary Information

### SUPPLEMENTARY FIGURES

**Supplementary Figure 1.**
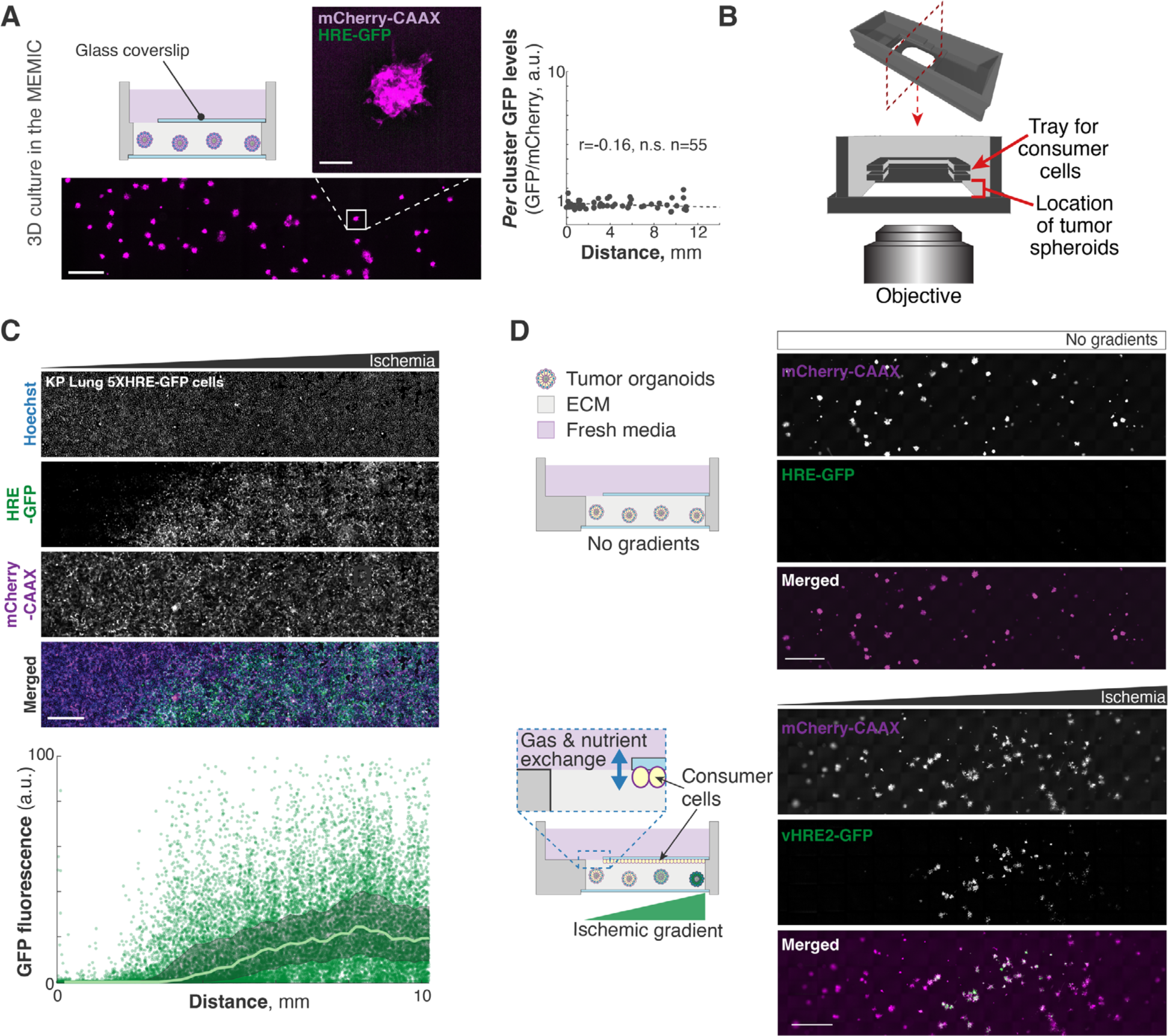
Consumer cells are required to establish ischemic gradients within the 3MIC. **A.** Spheroids grown in MEMICs fail to establish metabolic hypoxic gradients. Plot show the quantification of *per* spheroid GFP fluorescence (hypoxia reporter). Bars: 1000µm, and 100µm for insets. **B.** Schematic showing the design of the 3MIC. A coverslip with attached consumer cells is slid into the holder above embedded clusters to create a nearly enclosed chamber. The rest of the structure contains fresh media. **C.** Representative images of consumer cells expressing 5xHRE-GFP and mCherry-CAAX. Plot show the quantification of *per* spheroid GFP fluorescence (hypoxia reporter). Line: moving average. Bars: 1000µm **D.** Spheroids of C6 glioma cells stably expressing a different hypoxia reporter (vHRE2-GFP^73^). Bars: 1000µm.

**Supplementary Figure 2.**
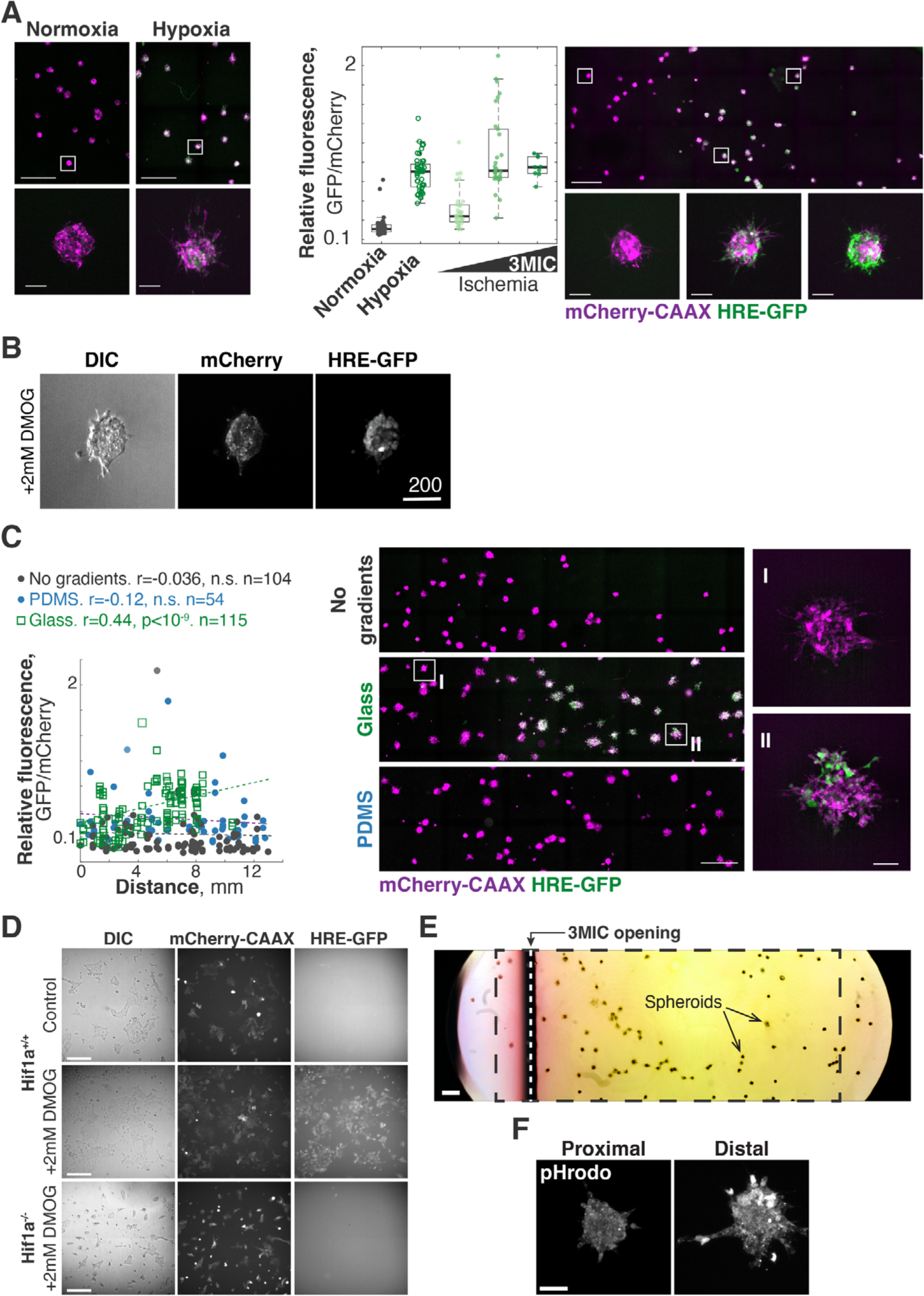
HIF1A and acidification of the extracellular space are critical for the increased invasion in ischemic cells. **A.** Lung KP spheroids were grown under normoxic, hypoxic and 3MIC conditions. Boxplots show GFP fluorescence (hypoxia) relative to mCherry-CAAX levels. Clusters in the 3MIC were grouped into three groups based on their distance from the opening. On the right, representative images of normoxic, hypoxic, and 3MIC conditions. Bars:1000µm,100µm for insets. **B.** Representative images of tumor spheroids treated with DMOG. Bars: 200µm. **C.** Lung KP spheroids expressing 5xHRE-GFP and mCherry-CAAX were grown without consumers, or in 3MICs enclosed in glass or PDMS. Points: *per* spheroid GFP (hypoxia) normalized to mCherry-CAAX levels. Lines: linear fit. Panels on the show representative images of spheroids grown under no consumer, glass, and PDMS 3MICs. Bars: 1000µm, 100µm for insets. **D.** 2D cultures of Hif1a^+/+^ and Hif1a^-/-^ cells treated or untreated with DMOG. These cells express HRE-GFP and mCherry-CAAX but as expected Hif1a^-/-^ cells do not express GFP in response to DMOG. Bars: 100µm. **E.** Color photo of a whole 3MIC with spheroids (see arrows) and consumer cells (not visible). Phenol red in the media shows a clear drop in extracellular pH. Black rectangle indicates the approximate area quantified in Fig. 2. Vertical white dashed line shows the opening of the 3MIC (shade below corresponds to the edge of the top coverslip). Bar: 500µm. **F.** Tumor spheroids were cultured in the 3MIC and labelled with an intracellular pH fluorescent probe (pHrodo). We did not observe significant differences in fluorescence between well-nurtured and ischemic spheroids.

**Supplementary Figure 3:**
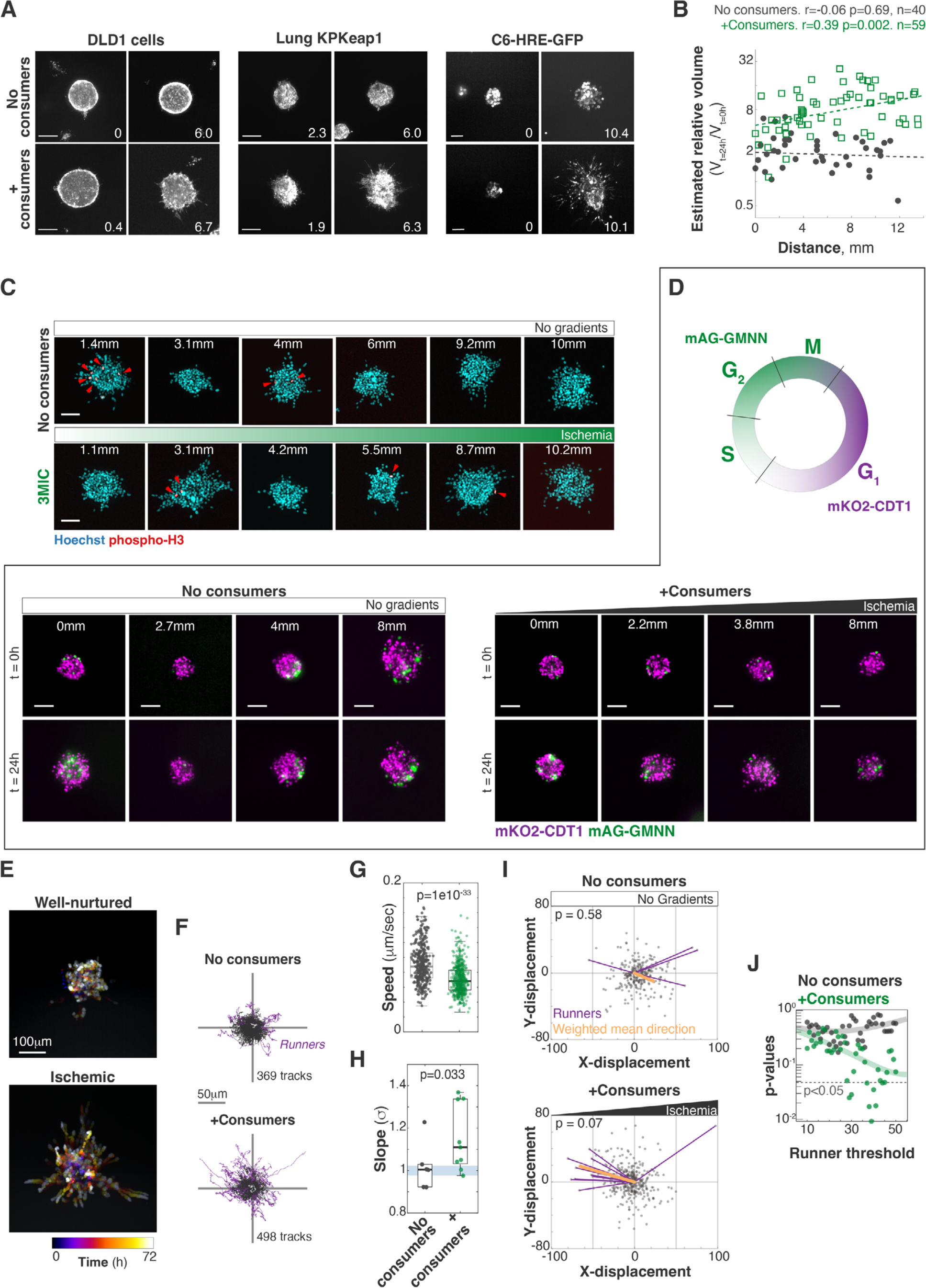
Ischemia increases cell motility and invasion. **A.** Representative images of spheroids of DLD1 cells (colorectal adenocarcinoma cells), KRAS^G12D^ p53-/- Keap1-/- cells (KPKeap1, lung adenocarcinoma) expressing 5xHRE-GFP and C6-hre-GFP cells (rat glioma) grown in 3MICs with or without consumers. Numbers in lower right corner denote the distance to the opening in mm. Bars: 100*μ*m. **B.** Quantification of estimated volume change of Lung KP spheroids. Points: area change of individual clusters, lines: linear fit. **C.** Phospho-H3 immunofluorescence staining of Lung KP spheroids in 3MICs. Red arrows: proliferating cells. Bars: 100µm. **D.** FUCCI-expressing lung KP clusters stably express mKO2-CDT1 in the G1 cell cycle phase or mAG-GMNN in other phases of the cell cycle. Clusters growing in the absence of ischemic gradients have many proliferating cells throughout the duration of the experiment, whereas ischemic clusters in the 3MIC have progressively lower numbers of proliferating cells, showing a higher portion of cells in G1. Bars: 100µm. **E.** Temporal projection of a representative ischemic cluster over a 72h time-lapse. **F.** Migratory tracks for all cells tracked in 3MICs and pooled according to the presence or absence of consumers. Tracks of cells that with a net displacement of 50μm or more (runners) are highlighted in magenta. **G.** Top: Instantaneous speeds of individual cells in the presence or absence of consumers. Bottom: MSD constants for cells moving in the presence or absence of consumers. Each dote is the MSD constant calculated from multiple tracks within one spheroid. **H.** Final position of each tracked cell centered to their origins. Magenta vectors highlight the displacement and direction of runner cells. Vector sum (orange) shows a small bias towards higher nutrient levels in the presence of consumers. However, this bias is only significant (Raleigh-Moore directional test) under some thresholds to define ‘runner cells’ (**I**).

**Supplementary Figure 4:**
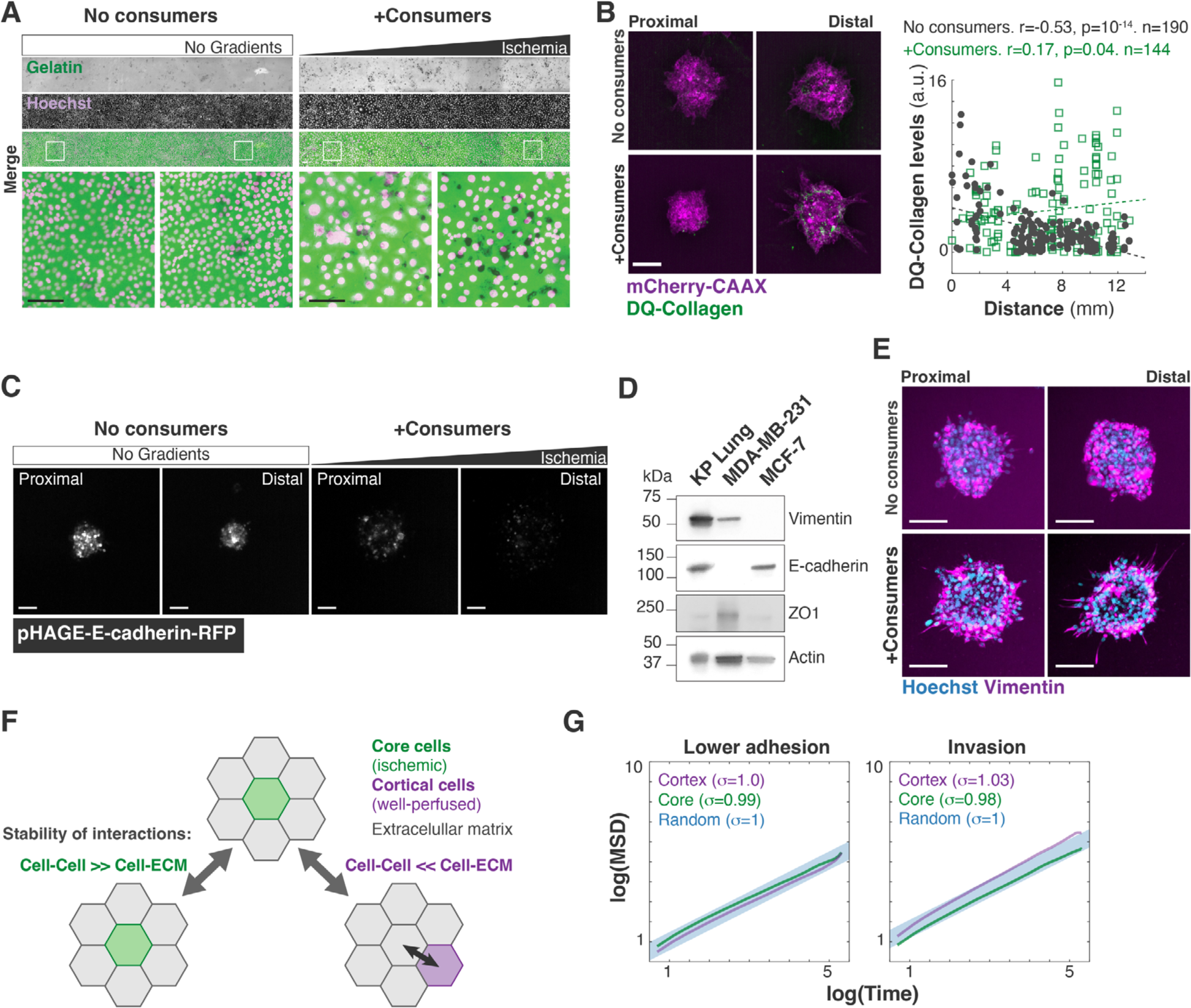
Ischemia enhances invasiveness of Lung KP spheroids. **A.** Lung KP cells were grown on 488-Gelatin coated coverslips and used as consumers. Increase in degradation of gelatin was observed with increase in distance from the opening in 3MIC, while no significant degradation was observed under no gradient conditions. Bars: 100µm. **B.** Right: representative images of Lung KP spheroids grown on DQ collagen. The distal ischemic spheroids increased their collagen/ECM cleavage. Bars: 100µm. Left: quantification of fluorescence intensity of DQ collagen showing a gradual increase in cleaved collagen with increase in distance of the spheroid from the opening. Points: DQ-collagen levels per cluster (GFP-fluorescence intensity normalized to mCherry-CAAX fluorescence). Dashed line: linear fit **C.** Lung KP tumor spheroids stably expressing pHAGE-E-cadherin-RFP (E-cadherin reporter). Bars: 100µm. **D.** Western blot analysis of EMT markers in Lung KP cells. **E.** Representative images of Lung KP spheroids grown in the 3MIC and stained for vimentin. Bars: 100µm. **F.** In the Cellular Potts model illustrating our model of ECM invasion. This model is mechanistically agnostic. It only says that in ischemic cells, the affinity to the ECM relative to cells in higher than in well nurtured cells. This simple rule increases the chance of a localization “swap” of ischemic cells into the ECM. **G.** Mean-squared displacement analysis for simulated well-nurtured cells and ischemic cells. Models included lower cell-cell adhesion in ischemia or increased invasion. The synergistic effects of both properties are shown in main figure 4.

**Supplementary Figure 5:**
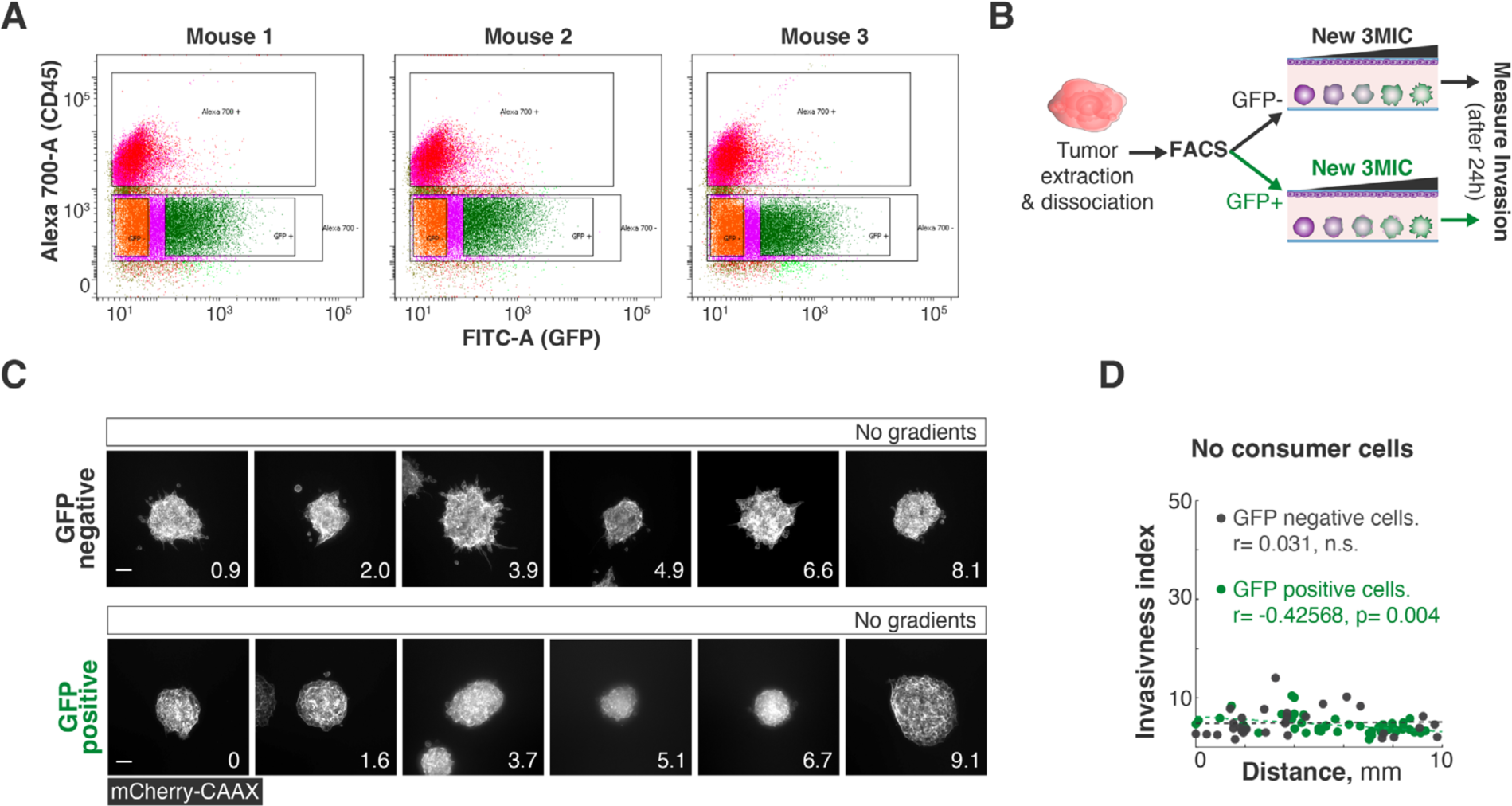
Ischemia-induced changes are phenotypic. **A.** 3x10^5^ Lung KP cells expressing 5xHRE-GFP were injected subcutaneously into the flank of syngenetic mice. Tumors were then dissociated into single cells which were then FACS-separated into GFP+ and GFP-populations. Plots show FACS data for tumor from three different mice. **B.** Experimental workflow: sorted cells were then grown in separate 3MICs with or without consumers. **C.** Representative images of spheroids culture in 3MICs with no gradients (no consumers). Bars: 100µm. **D.** In the absence of consumers, cells derived from hypoxic tumor regions did not show a significant increase of invasion or motility.

**Supplementary Figure 6:**
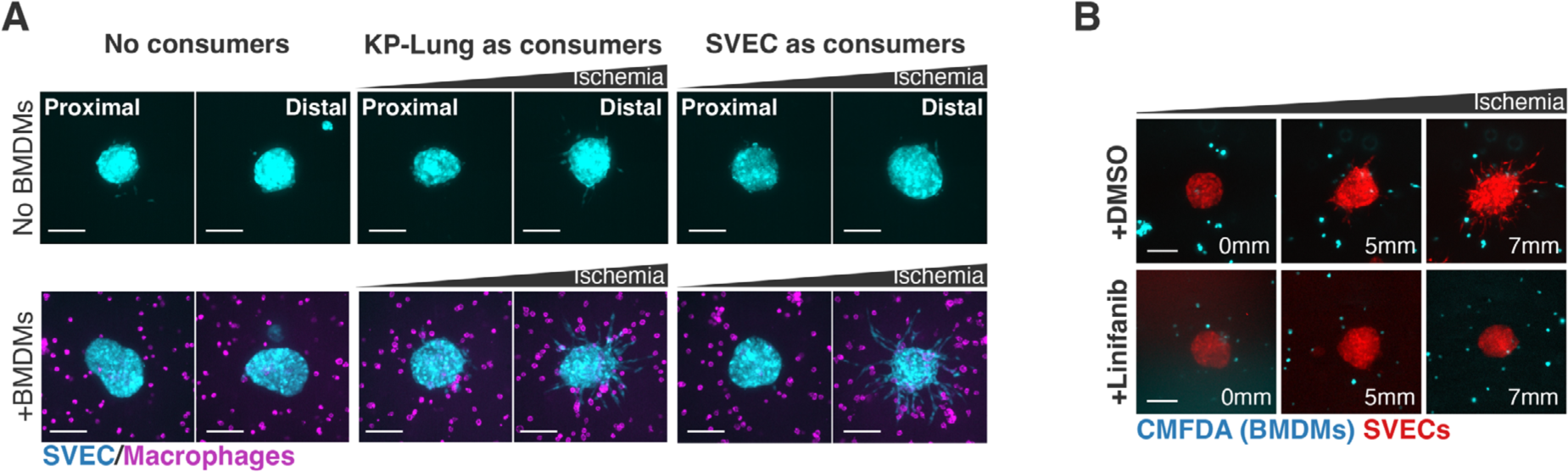
Ischemic macrophages induce sprouting of SVEC spheroids. **A.** Representative images of SVEC clusters co-cultured with macrophages in the absence of (no gradients) or presence of different cell types as consumers. Bars: 100µm. We did not observe a major effect of the consumer cell types but none replaced the pro-sprouting effects of macrophages. Images compare spheroids located 2mm or closer to the opening of the 3MIC (Proximal) or further than 8mm (Distal). **B.** In the presence of consumers (in this case also SVECs), BMDMs dramatically increase sprouting in SVEC clusters. This effect is abrogated by the VEGFA inhibitor Linifanib. Bars: 100µm.

**Supplementary Figure 7:**
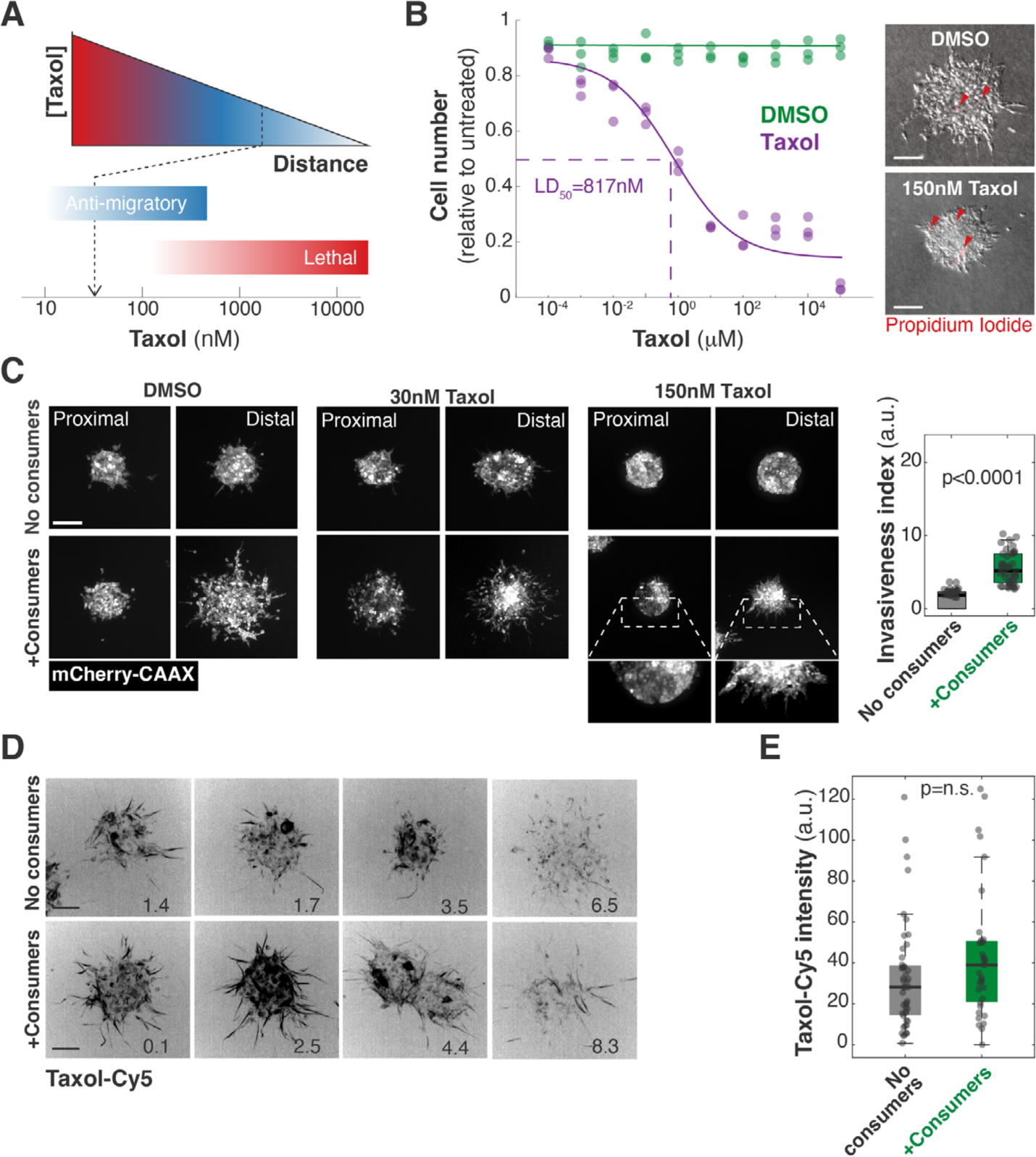
Ischemic spheroids are resistant to the anti-motility effects of Taxol. **A.** Schematic representation of lethal and sub-lethal as Taxol levels decrease with the distance to its source. We asked here is we can determine Taxol levels within 3MICs and test if those values have the same effects on ischemic cells and well-nurtured cells. We found that while about 30nM of Taxol inhibit the movements of well-nurtured cells, ischemic cells were not sensitive to these Taxol levels. **B.** Lung KP cells were treated with different doses of Taxol for 24 hours. Viability was determined by propidium iodide (PI) incorporation to determine the IC_50_ values. Representative images on the right illustrate that spheroids showed no significant differences in cell viability when spheroids were treated with doses of 150nM Taxol or lower. Arrowheads: Propidium positive non-viable cells. **C.** Representative images and quantification of Lung KP spheroids treated and untreated with 150nM Taxol. Distal spheroids can invade the ECM even when treated with Taxol but only in the presence of consumers (ischemia). Bars: 100µm. **D.** Representative images of Lung KP clusters grown in the 3MIC in the presence or absence of gradients and treated with a fluorescent Taxol analog (Taxol-Cy5). We used its fluorescent signal to estimate local levels of the drug in the 3MIC. Numbers in lower right corner denote the distance to the opening in mm. Bars: 100*μ*m. **E.** Boxplot showing Taxol-Cy5 levels in spheroids within 3MICs with or without consumer cells. Dots: *per* spheroid Taxol-Cy5 levels.

### SUPPLEMENTARY VIDEO LEGENDS

**Video S1.**
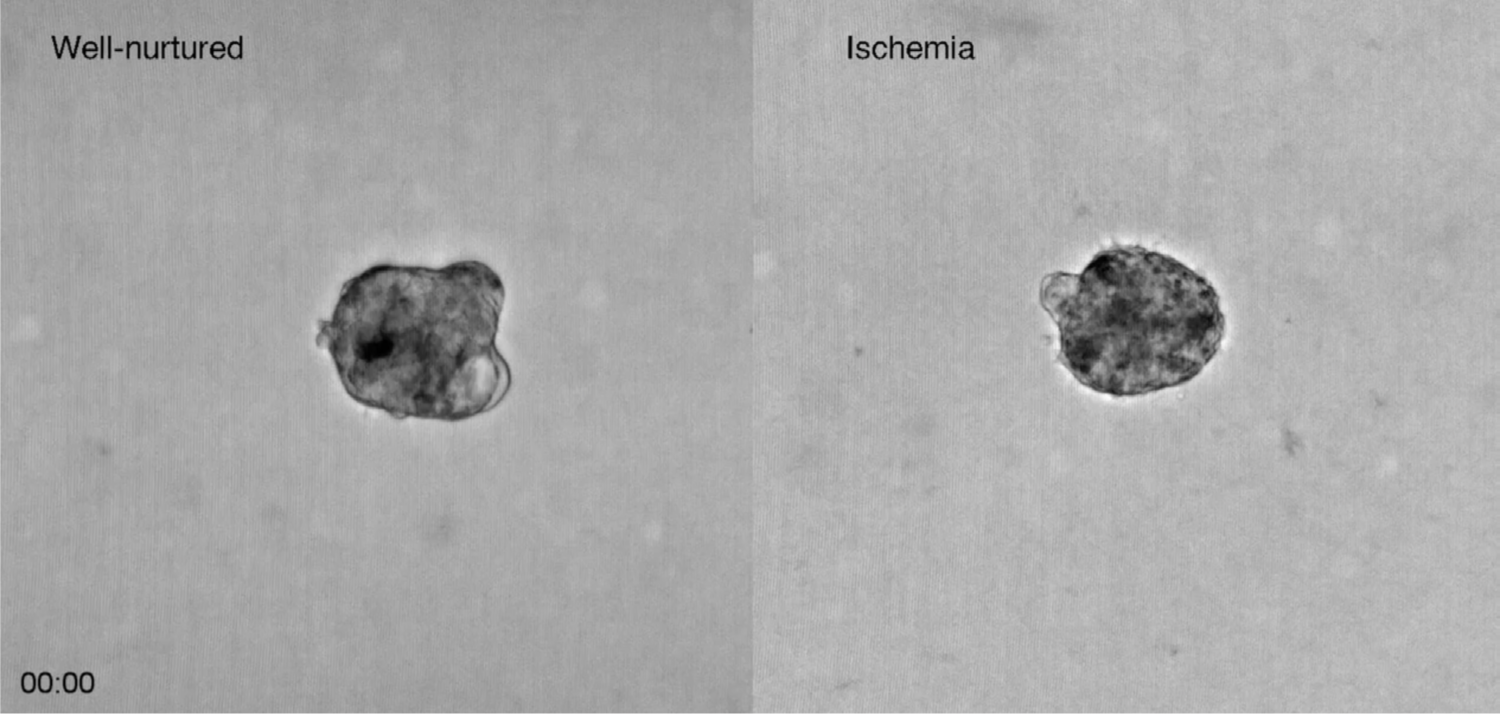
Lung KP spheroids grown in 3MIC. Left panel shows spheroid growing close to the opening of 3MIC (well-nourished) and right panel shows a spheroid growing in the ischemic region of 3MIC. Images were acquired every 15 mins for a period of 72 hours.

**Video S2.**
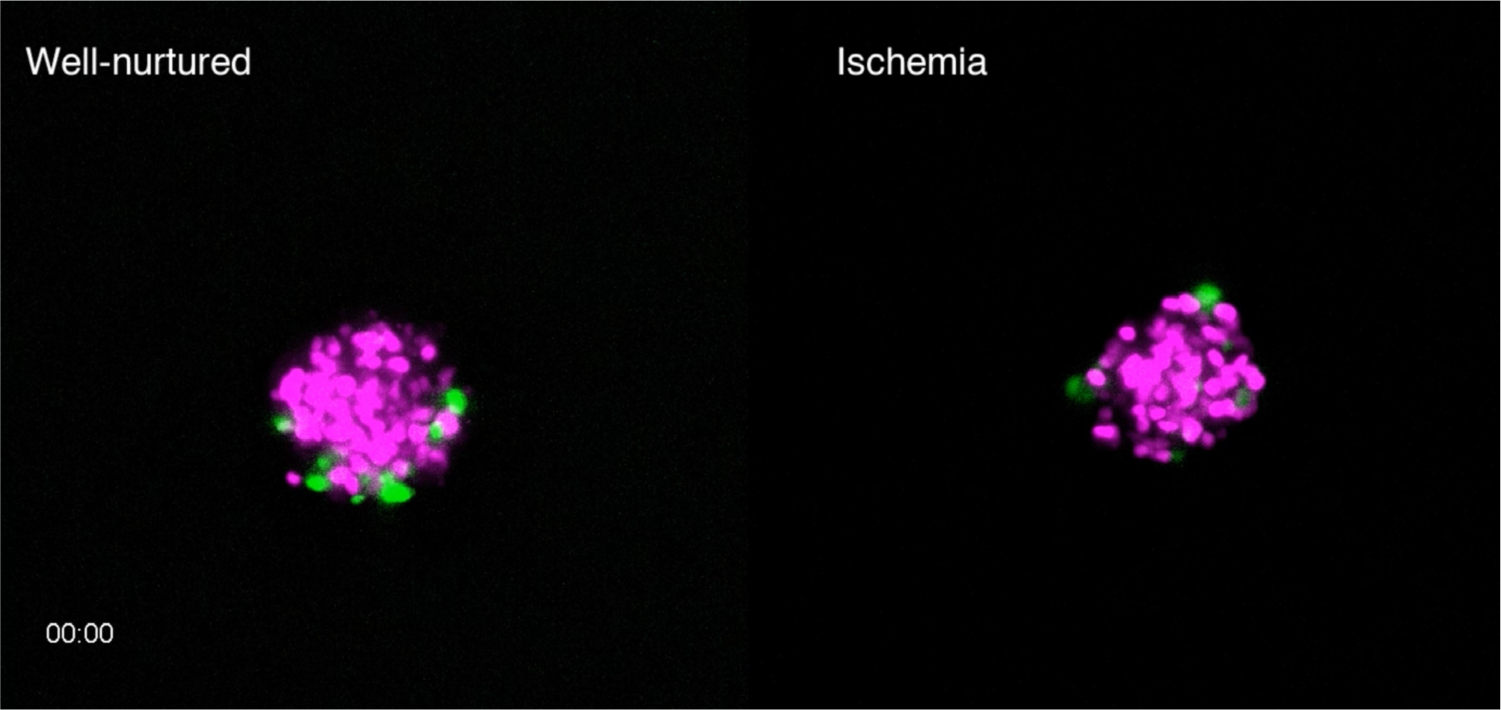
Lung KP spheroids stably expressing the pBOB-EF1-FastFUCCI-Puro construct were grown in 3MIC. Movie shows the cell-cycle progression of the cells in the spheroids growing in well-nourished regions (left panel) and ischemic regions (right panel).

**Video S3.**
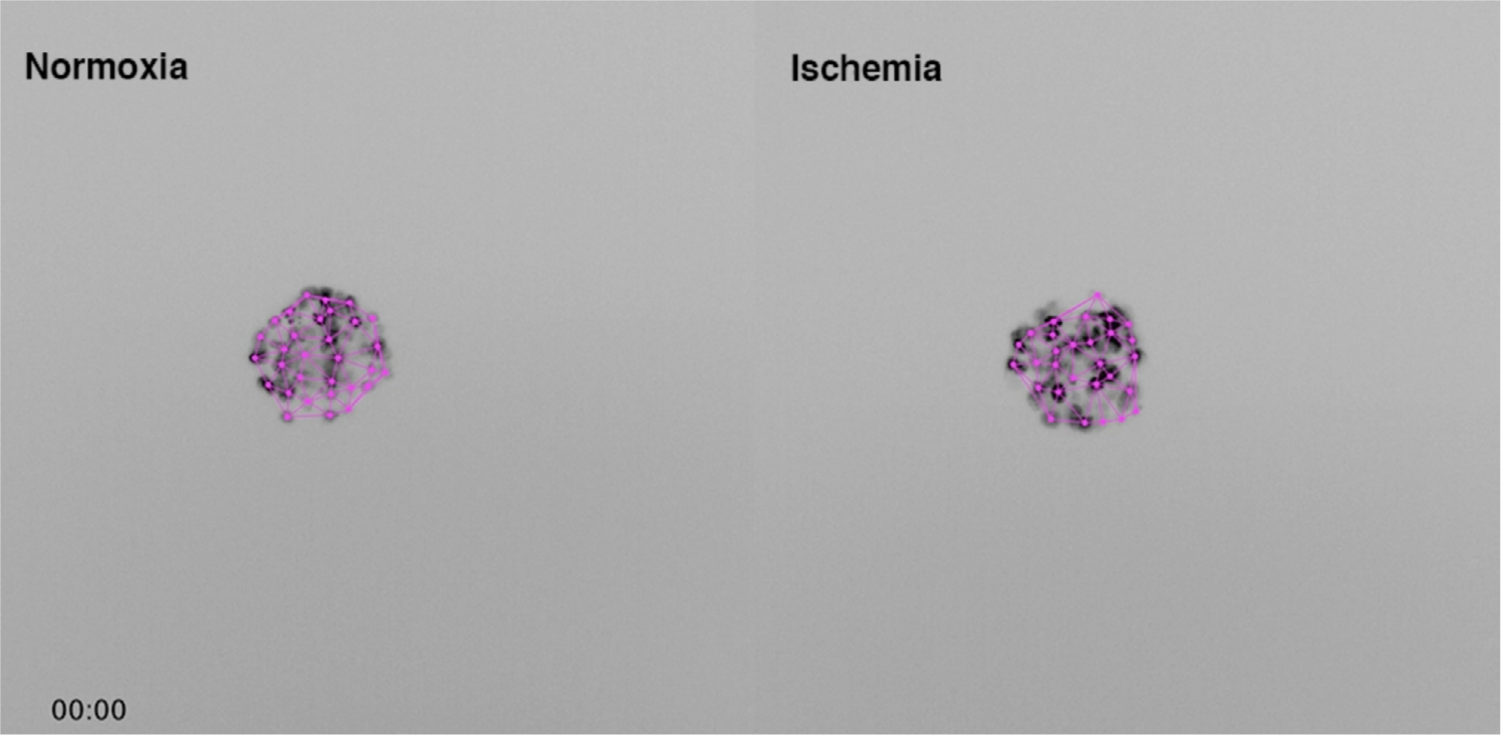
Movie depicting the decrease in cell density (or increase of area in between cells). Cells are labelled with H2B-YFP (black). Purple lines show results from Delaunay triangulation algorithm.

**Video S4.**
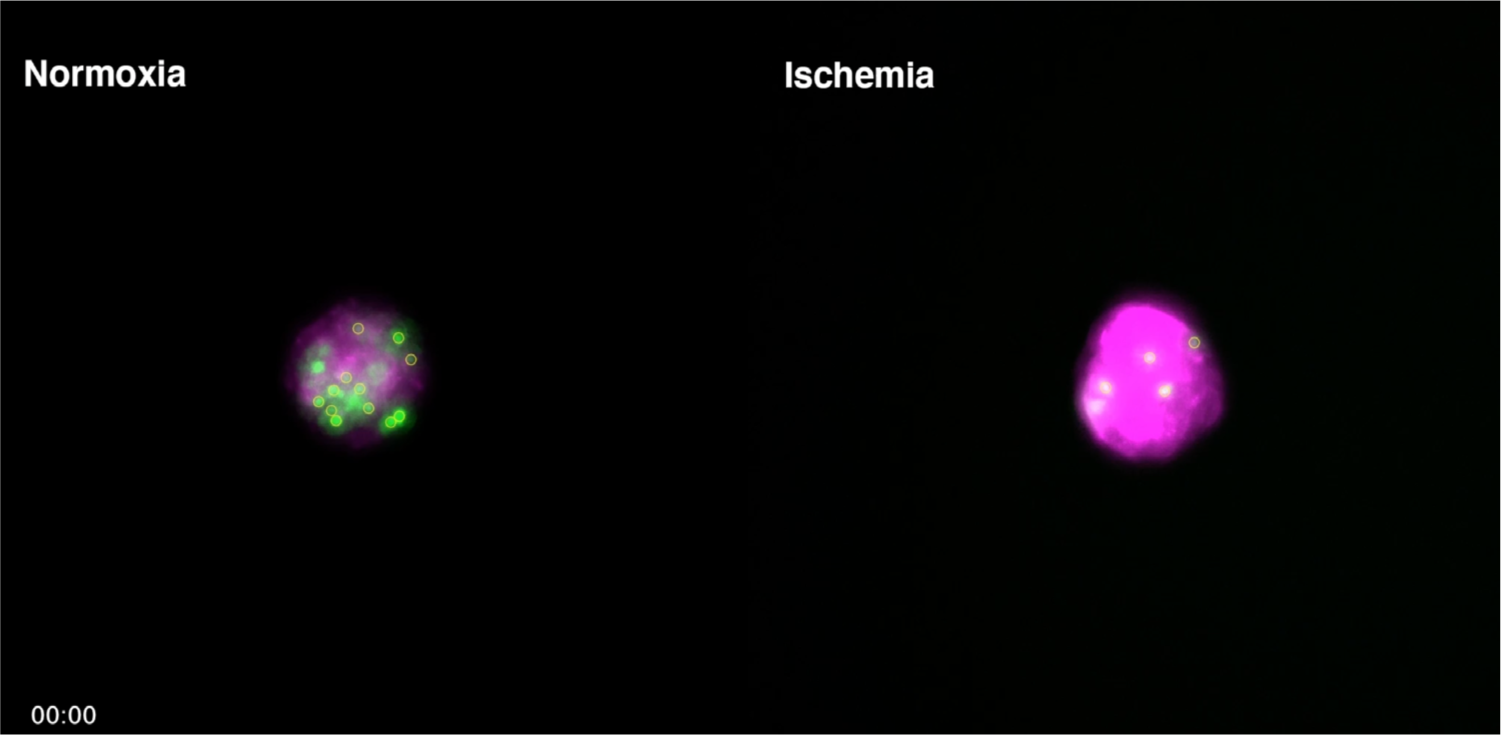
Chimeric Lung KP spheroids of cells expressing H2B-YFP (green) or mCherry-CAAX (magenta) mixed at a 1:4 ratio. Time-lapse video shows the migration tracks of H2B-YFP positive cells in spheroids grown in well-nurtured and ischemic regions of 3MIC.

**Video S5.**
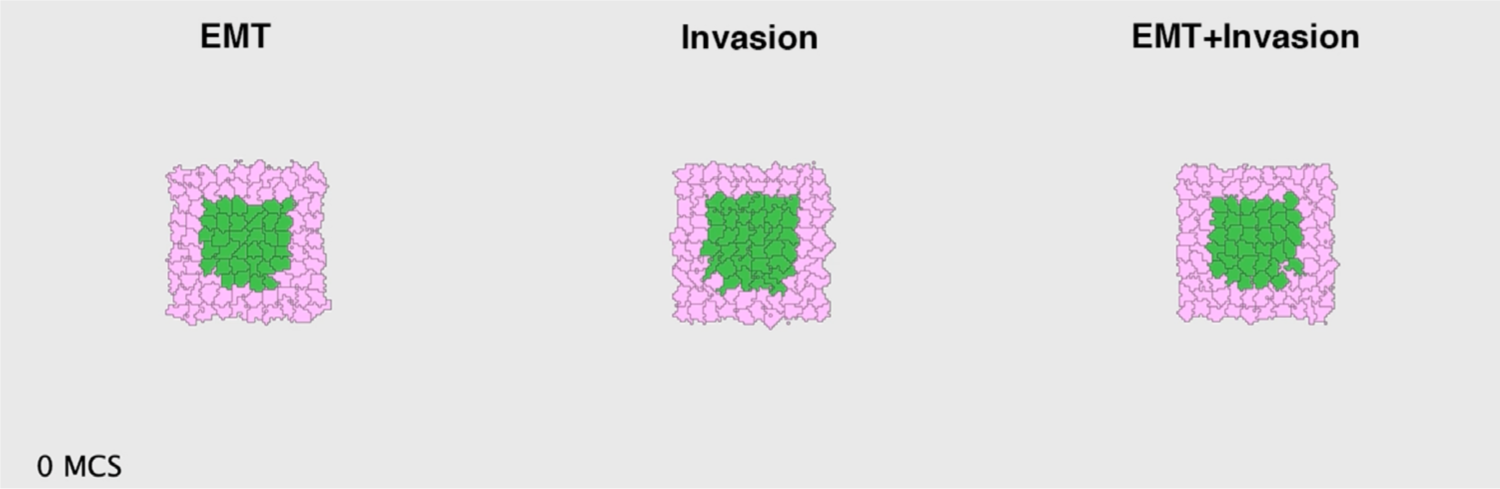
Movie depicting the evolution of three Cellular Potts Models simulating the movements of cells arranged as a core of ‘ischemic’ cells surrounded by cortical, well-nurtured cells. In these models, core cells are assigned a reduced epithelial cell adhesion, increased ECM invasion, or the combination of both using. MCS: Monte Carlo Step (time unit).

**Video S6.**
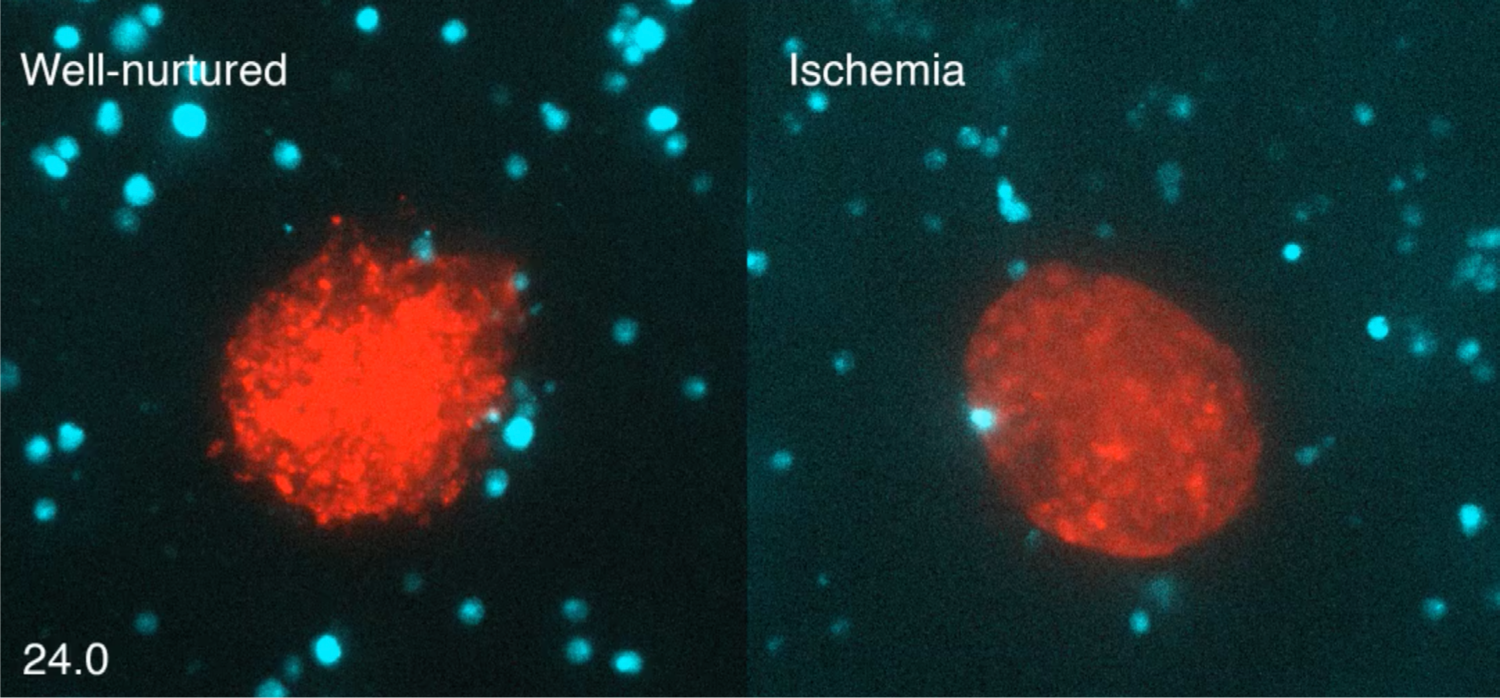
Time lapse of SVEC spheroids (expressing mCherry, red), growing in the 3MIC with BMDMs (labelled with CMFDA, cyan) embedded in the ECM. Images were acquired from 24 hours every 30 mins for a period of 72 hours.

**Video S7.**
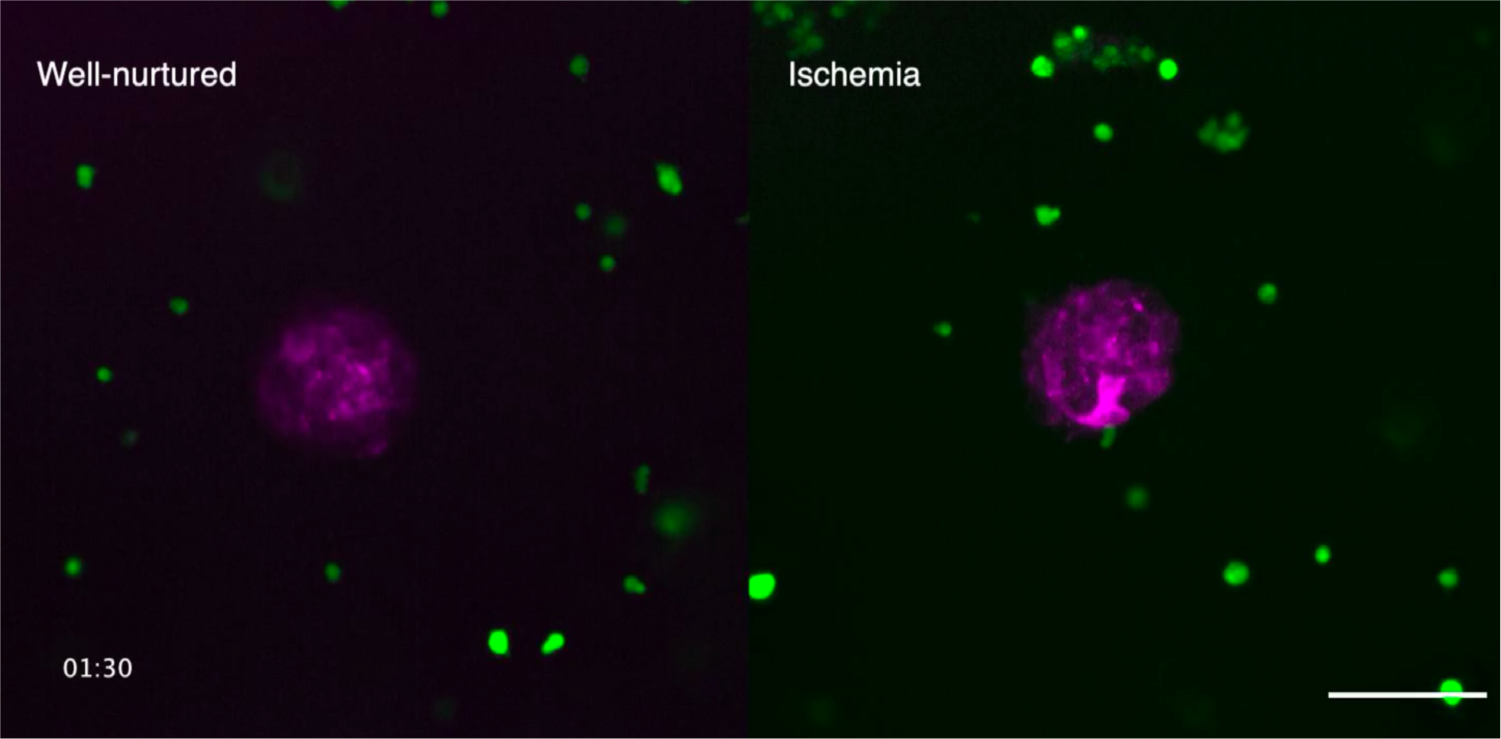
Time lapse of Lung KP (expressing mCherry-CAAX, magenta) spheroids growing in 3MIC in the 3MIC with BMDMs (labelled with CMFDA, green) embedded in the ECM. Images were acquired every 30 mins for a period of 72 hours.

**Video S8.**
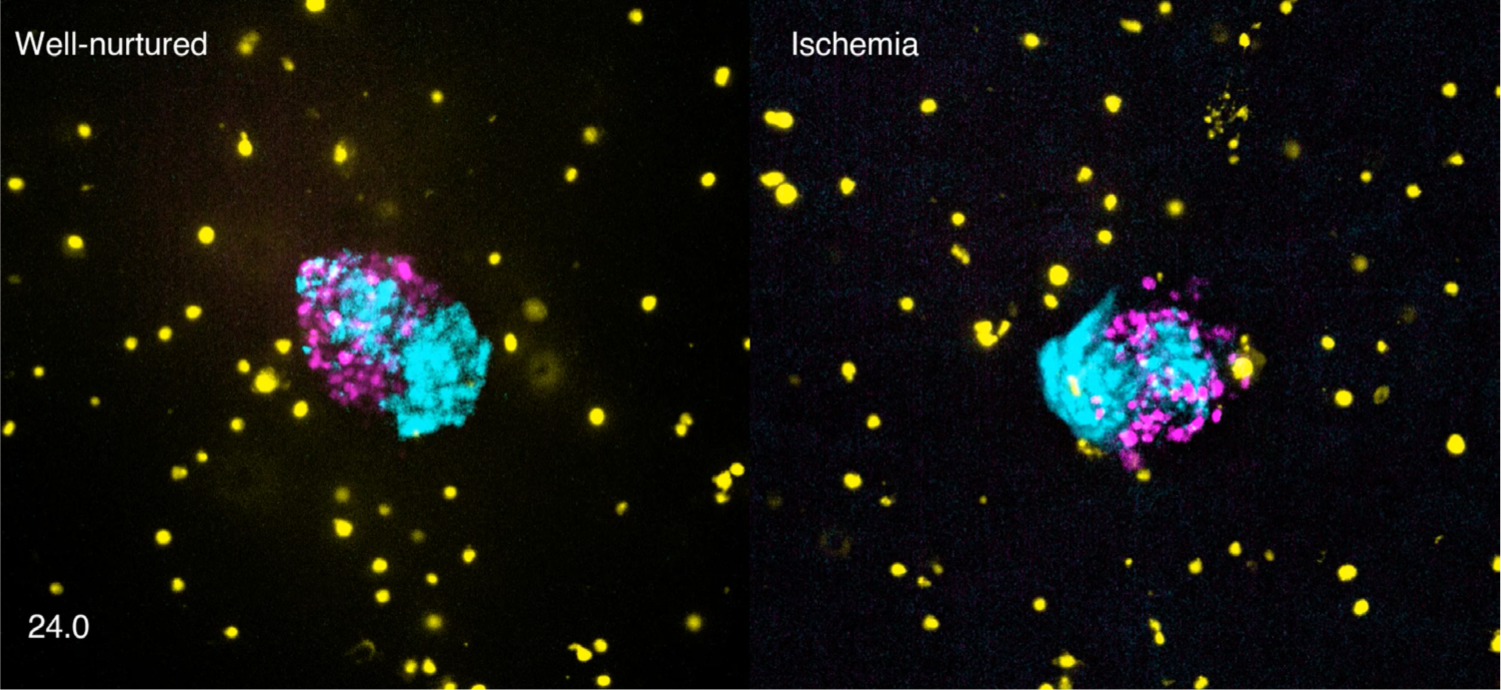
Time lapse of spheroids formed by Lung KP cells (magenta) and SVECs (cyan), growing in the 3MIC with BMDMs (yellow) embedded in the ECM. Images were acquired from 24 hours every 30 mins for a period of 66 hours.

### SUPPLEMENTARY TABLES

**Table S1.** CPM parameters for different conditions. *J*_*C*1,*C*1_ is the surface energy between to cells or a cell and the ECM. Higher surface energies lead to lower affinity. C1: core cells. C2: cortical cells. ECM: Substrate (extracellular matrix).

| Condition | $J_{C1,C1}$ | $J_{C1,C2}$ | $J_{C2,C2}$ | $J_{C1,ECM}$ | $J_{C2,ECM}$ |
| --- | --- | --- | --- | --- | --- |
| Increased Invasion | 16 | 16 | 16 | 2 | 11 |
| Decreased epithelial adhesion | 24 | 16 | 8 | 11 | 11 |
| Both | 24 | 16 | 8 | 2 | 11 |

**Table S2.** Additional CPM parameters.

| Parameter | Value | Units |
| --- | --- | --- |
| Temperature | 37 | °C |
| Population size | 100 | cells |
| Target cell size | 40 | pixels |
| Dimensions | 200x200 | pixels |
| Boundary conditions | Periodic | N/A |

